# Roles of SNORD115 and SNORD116 ncRNA clusters in neuronal differentiation

**DOI:** 10.1101/2023.10.24.563766

**Authors:** Aleksandra Helwak, Tomasz Turowski, Christos Spanos, David Tollervey

**Affiliations:** Wellcome Centre for Cell Biology, The University of Edinburgh, Edinburgh, Scotland; Institute of Biochemistry and Biophysics PAS, Warszawa, Poland

## Abstract

Prader-Willi syndrome shows features linked to brain development and hypothalamus-related endocrine abnormalities. The smallest clinical deletions fall within the large (∼650Kb) SNHG14 gene, removing 29 consecutive introns that each generate SNORD116. SNHG14 also includes 48 tandem introns encoding SNORD115 and generates multiple, extended snoRNA-related species. SNORD115 and SNORD116 resemble box C/D small nucleolar RNAs (snoRNAs) but lack known targets. Both snoRNAs strongly accumulated during neuronal differentiation. SNORD116 accumulation apparently reflected stabilization, potentially linked to the appearance of FBLL1, a homologue of the ubiquitous snoRNA-associated protein Fibrillarin (FBL). In contrast, SNORD115 was selectively transcribed, apparently due to regulated termination. For functional characterization we created cell lines lacking only the expressed, paternal, SNORD115 or SNORD116 cluster. Analyses during neuronal development indicated changes in RNA stability and protein synthesis. Altered mRNAs included *MAGEL2*, mutation of which causes the PWS-like disorder Schaaf-Yang syndrome. Comparison of SNORD115 and SNORD116 mutants indicated overlapping or interacting functions. Most changes in mRNA and protein abundance appeared relatively late in development, with roles including cytoskeleton formation, extracellular matrix, neuronal arborization. Comparison with human embryonic midbrain development suggested enhanced progression in neuronal development in the snoRNA mutants. Subtle impairment of relative neuronal maturation during development, might generate the clinical phenotypes.

## INTRODUCTION

Prader-Willi syndrome (PWS) is a paradigm of neurodevelopmental disorders with a frequency of ∼1:20,000 (Bieth *et al*, 2015; Bochukova *et al*, 2018; Bortolin-Cavaille & Cavaille, 2012; Cavaille *et al*, 2002) caused by deletions in chromosome 15 (15q11.2-q13) (Fig. 1). The region is “imprinted” with different expression patterns for the paternal and maternal chromosomes. PWS specifically results from the lack of expression of genes from the paternal chromosome, due to deletion, uniparental disomy or imprinting center defects. Deletions in the maternal chromosome cause a different neurological disease, Angelmann syndrome. This has been linked to loss of *UBE3A* which encodes a ubiquitin ligase ((Wolter *et al*, 2020) and references therein). Deletions causing PWS typically remove large genomic regions, however, disease-linked microdeletions have also been identified (Bieth *et al*., 2015; Duker *et al*, 2010; Tan *et al*, 2020). The smallest remove a ∼70 Kb region of *SNHG14* (snoRNA host gene 14), in which 29 tandem introns each encode the small nucleolar RNA (snoRNA) SNORD116 (Fig. 1) (Cavaille *et al*, 2000; Cavaille *et al*., 2002; Duker *et al*., 2010).

**Figure 1.**
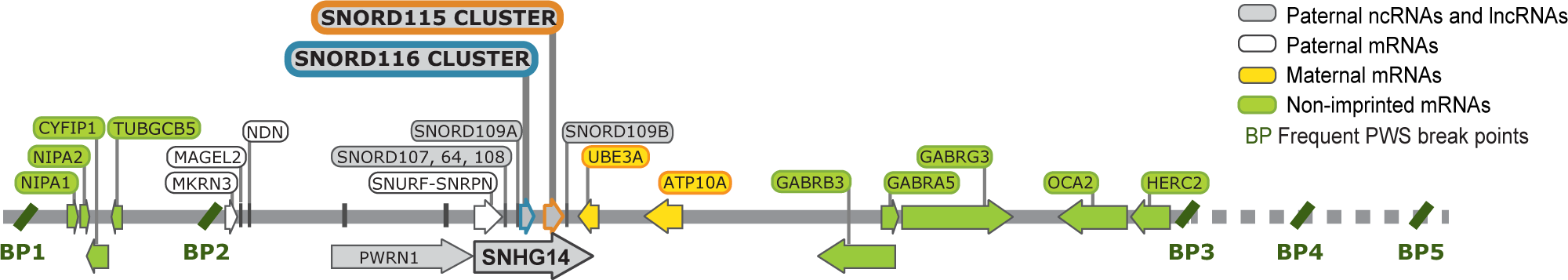
Scheme of PWS locus.

*SNHG14* generates a very long non-protein coding RNA (lncRNA) with a predicted primary transcript around 600 Kb in length including 145 annotated introns (Runte *et al*, 2001). It is processed into multiple ncRNAs; including mature snoRNAs, extended snoRNA-related ncRNA species (SPA-lncRNAs and sno-lncRNAs), and alternatively spliced versions of the *SNHG14* exons ((Sledziowska *et al*, 2023; Yin *et al*, 2012) reviewed in (Ariyanfar & Good, 2022)). Multiple, non-identical versions of SNORD116, are encoded by 29 tandem introns of *SNHG14* and excised following splicing. Adjacent to the SNORD116 region a further 48 tandem introns encode another snoRNA-like species, SNORD115. However, differences in the spatiotemporal expression profiles of SNORD115 and SNORD116 have been reported (Landers *et al*, 2004; Vitali *et al*, 2010), potentially reflecting a boundary conferring tissue-specific expression of the *SNORD115* and *UBE3A-ATS* regions (Hsiao *et al*, 2019; Martins-Taylor *et al*, 2014). Splicing of the snoRNA-containing introns in mouse transgenes was inhibited by depletion of the neuronal splicing factor RBFOX3 (NeuN) (Coulson *et al*, 2018). In humans, most snoRNA species are encoded within introns of mRNAs or long non-protein coding RNAs. In analyzed cases the mature snoRNAs are generated by 5’ and 3’ exonuclease degradation of the excised intron following splicing and debranching of the intron lariat. Progression of the exonucleases is likely blocked by snoRNA assembly with proteins into stable, small nucleolar ribonucleoprotein (snoRNP) particles, since loss of these proteins in yeast prevents snoRNA accumulation.

PWS individuals show a range of developmental and neurological deficits. Perhaps most notable is hyperphagia, which leads to potentially life-threating over-eating. This has been linked to altered gene expression in the hypothalamus, where hunger is regulated (Bochukova *et al*., 2018; Polex-Wolf *et al*, 2018; Tauber *et al*, 2014). Human tissue distribution data confirmed high SNORD116 levels in multiple brain regions including, but not limited to, the hypothalamus These findings suggest direct roles for the snoRNA in gene expression leading to regulated feeding.

SNORD116 and SNORD115 species resemble box C/D class snoRNAs, which have characteristic structural features and bind a set of four, highly conserved proteins; Fibrillarin (FBL), NOP56, NOP58 and SNU13, (NPHX, 15.5K) (Fig. 4A). All characterized snoRNAs function through base-pairing with target RNAs, most commonly directing site-specific modification in rRNAs or other small stable ncRNAs, while some are required for pre-rRNA processing. Box C/D snoRNAs generally form extended base-paired interactions that precisely target the 2’-hyrodroxyl residue on the nucleotide located 5 base-pairs from the box D motif in the snoRNA. This directs nucleotide-specific 2’-*O*-methylation of the ribose group by the snoRNA-associated methyltransferase FBL. This specificity allows target sites for many snoRNAs in rRNA and stable ncRNAs to be precited with considerable confidence. However, no relevant targets are known for SNORD116 (Cavaillé *et al*, 2000). Potential targets in mRNAs have been predicted (Baldini *et al*, 2022), and reporter constructs indicate that SNORD116 expression can stabilize the NHLH2 mRNA (Burnett *et al*, 2017; Kocher *et al*, 2021)

Most snoRNAs are ubiquitously expressed, but ∼200 including SNORD116 and SNORD115 were reported to show brain-enriched expression (Cavaillé *et al*., 2000). Most of these are described as “orphans” since, like SNORD116 and SNORD115, they lack evident base-complementarity to rRNA or other targets. Loss of SNORD115 was previously reported to impair specific pre-mRNA splicing and editing events on mRNAs. These include that encoding neuronal serotonin receptor 2C (5HT2C) (Falaleeva *et al*, 2015; Kishore & Stamm, 2006), although this finding was questioned (Hebras *et al*, 2020). Changes in mRNA levels have been reported in human PWS-derived cells, brain samples and SH-SY5Y cells, but with limited consistency ((Baldini *et al*., 2022; Bochukova *et al*., 2018; Burnett *et al*., 2017; Powell *et al*, 2013); reviewed in (Bochukova, 2021)). Overexpression of SNORD116 and SNORD115 in non-neuronal cells also altered the abundance of many RNAs, with apparent interactions when co-expressed (Falaleeva *et al*., 2015). In addition, extended forms of the snoRNAs have been proposed to sequester specific RNA-binding proteins, including the pre-mRNA spicing factor Rbfox2 (Wu *et al*, 2016; Yin *et al*., 2012). The *SNHG14* host gene is widely expressed, with brain-enrichment, and other ncRNA products could also be disease-related (reviewed in (Ariyanfar & Good, 2022)). Despite these findings, the mechanistic basis for links between non-coding RNAs originating from the *SNHG14* locus and PWS remain unclear.

In a neuronal model, we tested mechanisms by which RNAs encoded by SNORD115 and SNORD116 clusters might alter gene expression, including: (1) impaired ribosome synthesis (analysis of pre-rRNA processing); (2) changes in RNA abundance (RNA-sequencing); (3) changed alternative splicing (RNA-sequencing); (4) translation efficiency (comparing transcriptome [RNA-sequencing] to proteome [mass spectrometry]); (5) TF sequestration (bioinformatics of key changes in gene expression). Substantial changes were observed in gene expression, at the levels of RNA abundance and predicted translation. These could not be linked to direct snoRNA base-pairing, but the transcriptome profile of cells depleted of SNORD116 indicated that the effect is to advance developmental progression.

## RESULTS

### Regulated expression of ncRNAs from the *SNHG14* locus

Consistent with their reported brain-enriched expression (Cavaillé *et al*., 2000) we saw no SNORD115 or SNORD116 in HEK293 cells. However, we detected expression in human Lund human mesencephalic (LUHMES) cells following differentiation to neurons. LUHMES are an embryonic, mid-brain derived human cell line that can be induced to synchronously differentiate into polarized dopaminergic neurons (Lauter *et al*, 2020; Smirnova *et al*, 2016) (Fig. S1A). Neuronal markers are expressed by day 6 of differentiation; including β3-tubulin (TUBB3), postsynaptic density protein 95 (PSD95, postsynaptic marker), kinesin 17 (KIF17, neuronal transport marker) and tyrosine hydroxylase (TH) (Lauter *et al*., 2020; Smirnova *et al*., 2016). This high synchrony was of particular importance for biochemical analyses during differentiation time courses. We tested snoRNA expression over differentiation time course up to day 15, after which LUHMES cells become sensitive to detaching from the dish. In cycling, pre-neuronal cells (day 0; D00) SNORD115 was not detectable by northern hybridization and SNORD116 was at low abundance (Fig. 2A, S2B). Both rose rapidly during differentiation, with a plateau for SNORD116 between day 6 (D06) and day 10 (D10), whereas SNORD115 steadily increased up to day 15 (D15). This relatively late expression of SNORD115 and SNORD116 suggested their importance during later stages of neuronal maturation. RNA sequencing (RNAseq) was therefore performed on undifferentiated cells and during differentiation at D06, D10 and D15. Note that the RNAseq approach used does not detect mature snoRNAs due to their short lengths. LUHMES cells are reported to be diploid (Shah *et al*, 2016). Normal karyotyping cannot exclude specific duplication in the PWS region. However, RNAseq confirms deletions within the expressed, paternal *SNHG14* gene, whereas maternal expression of the non-overlapping, convergent UBE3A gene is unaffected.

**Figure 2.**
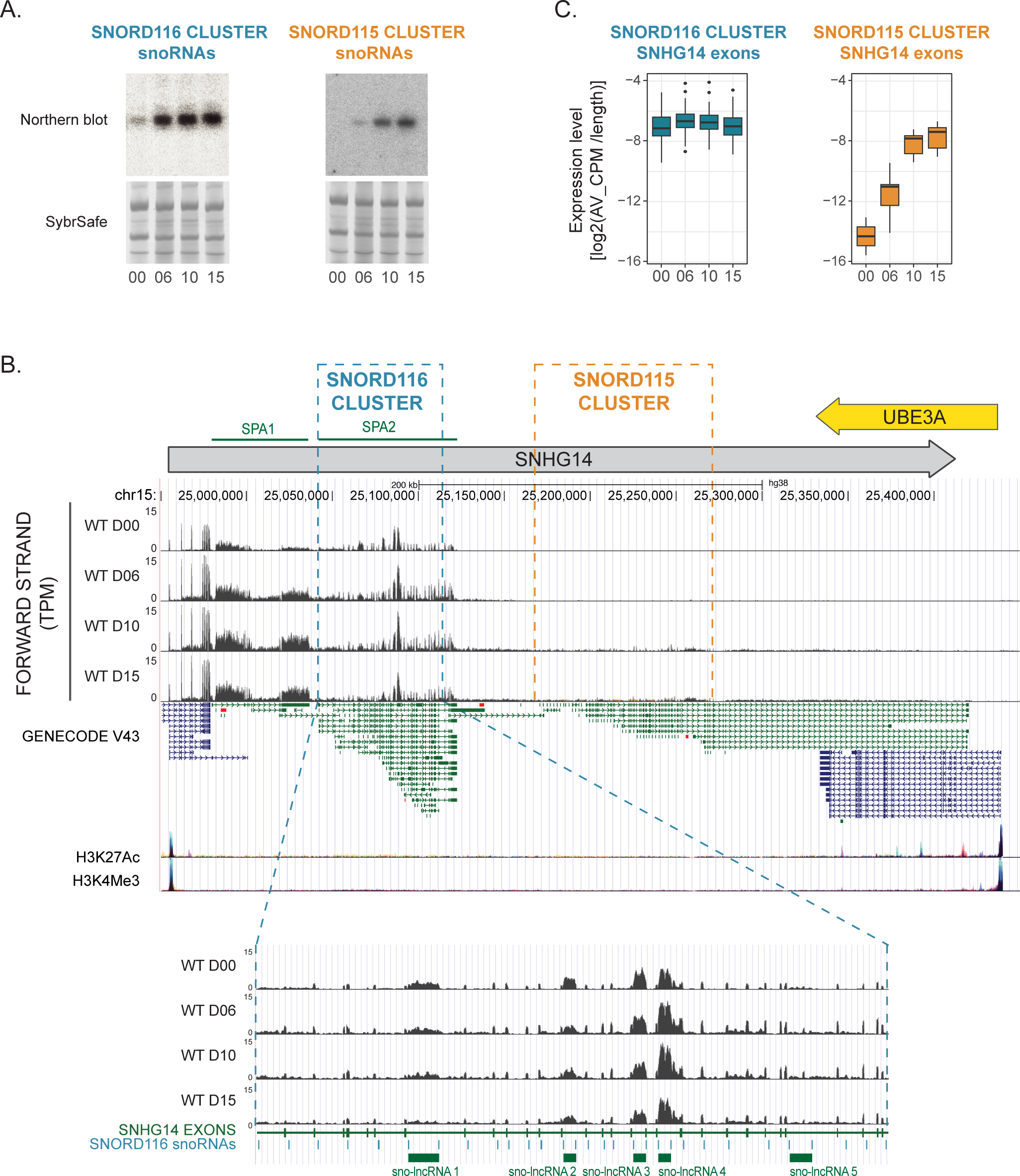
Changes in expression of ncRNAs from SNHG14 gene during neuronal differentiation. **A**: Expression of SNORD115 and SNORD116 snoRNAs in LUHMES cells upon differentiation. Northern blot and corresponding SybrSafe stained fragment of the gel below. **B**: UCSC Genome Browser view of the expression from the SNHG14 gene at day 0, 6, 10 and 15 of differentiation. GENECODE 43 track displays basic gene set with splice variants, mRNAs (blue) and non-coding RNAs (green). Layered H3K27Ac and H3K4Me3 tracks display data on histone modifications from ENCODE project, associated with the enhancer/regulatory regions and promoters, correspondingly. SNORD116 and SNORD115 cluster regions within the SNHG14 gene are marked with boxes. SPA and sno-lncRNAs ncRNAs, previously described but not included in the GENECODE track, are marked at the top and bottom of the figure. **C**: Difference in expression profiles of SNHG14 lncRNA exons, overlapping either SNORD116 or SNORD115 cluster. Expression levels (CPM) for individual exons are length normalized.

RNAseq data was initially characterized for expression of ncRNAs originating from the *SNHG14* gene (Fig. 2B). SNHG14 RNA abundance over the region surrounding the *SNORD116* cluster was almost unaltered during differentiation. Accumulation of the exons was substantially greater than for introns, which are normally rapidly degraded following debranching, consistent with correct splicing of introns encoding SNORD116 throughout differentiation. In contrast, the region surrounding the SNORD115 cluster showed low expression at D00. This increased during differentiation, but remained substantially lower than around *SNORD116* even at D15. To better characterize the changes in expression, reads mapping to exon sequences were compared (Fig. 2C). This confirmed that transcripts around *SNORD115* markedly increased during differentiation, while those around *SNORD116* and at sites further 5’ were essentially unchanged. Extended snoRNA-related RNAs SPA1 and sno-lncRNAs 1-4 (Sledziowska *et al*., 2023; Wu *et al*., 2016; Yin *et al*., 2012) were readily detected (Fig. 2B), but showed only modest changes during differentiation. Other reported ncRNAs, SPA2 and sno-lncRNA 5, were not clearly identified.

The sequence data showed a clear drop between the SNORD116 and 115 clusters, whereas *SNHG14* was previously described as a single ∼596 Kb transcription unit. To investigate this, we inspected mapping data for histone H3 lysine 4 trimethylation (H3K4me3), characteristic of RNAPII transcription initiation sites, and H3K27Ac, characteristic of regulatory regions (see; www.ncbi.nlm.nih.gov/gene/104472715 and UCSC genome browser ENCODE regulation tracks) (Kent *et al*, 2002). H3K4me3 peaks, and accompanying H3K27Ac peaks, were found at the predicted transcription start sites for *SNHG14* and the flanking *UBE3A* protein coding gene (Fig. 2B). There was no indication of initiation between the *SNORD116* and *SNORD115* clusters. We therefore predict that the apparent extension of the *SNHG14* transcripts into the region surrounding the *SNORD115* cluster reflects regulated read-through of a termination site located 3’ to *SNORD116,* as previously proposed (Vitali *et al*., 2010). Consistent with this, GENECODE V43 (Fig. 2B) indicates processing of *SNHG14* transcripts into multiple alternatively-spliced versions (in non-neuronal cells), covering *SNORD116* or *SNORD115* but not overlapping both clusters.

We conclude that the region encoding *SNORD116* is well transcribed in undifferentiated cells. The accumulation of exon regions relative to introns indicates that splicing is functional, suggesting that the failure in mature *SNORD116* accumulation reflects instability. We speculate that impaired assembly with snoRNP proteins allows degradation of the snoRNA sequence along with the excised intron within which it is embedded. At the same time there is abundant expression of SNORD116 containing sno-lncRNAs, indicating that those two kinds of ncRNAs undergo distinct, possibly competing processing pathways. In contrast, *SNORD115* is poorly expressed in undifferentiated cells, with increased transcription readthrough into this region during differentiation.

### Transcriptional changes during differentiation of LUHMES cells

LUHMES cells are frequently used as a model of dopaminergic neurons. Changes in the transcriptome (Lauter *et al*., 2020) and proteome (Tüshaus *et al*, 2021) were reported up to day 6 of neuronal differentiation, at which point they were considered mature. In agreement with that, in our data most changes to the transcriptome occur between D00 and D06 of LUHMES differentiation (Fig. 3A). However, we observe that about 1% of genes that change expression during differentiation, showed changes between D10 and D15, when snoRNA-related effects appeared more likely.

**Figure 3.**
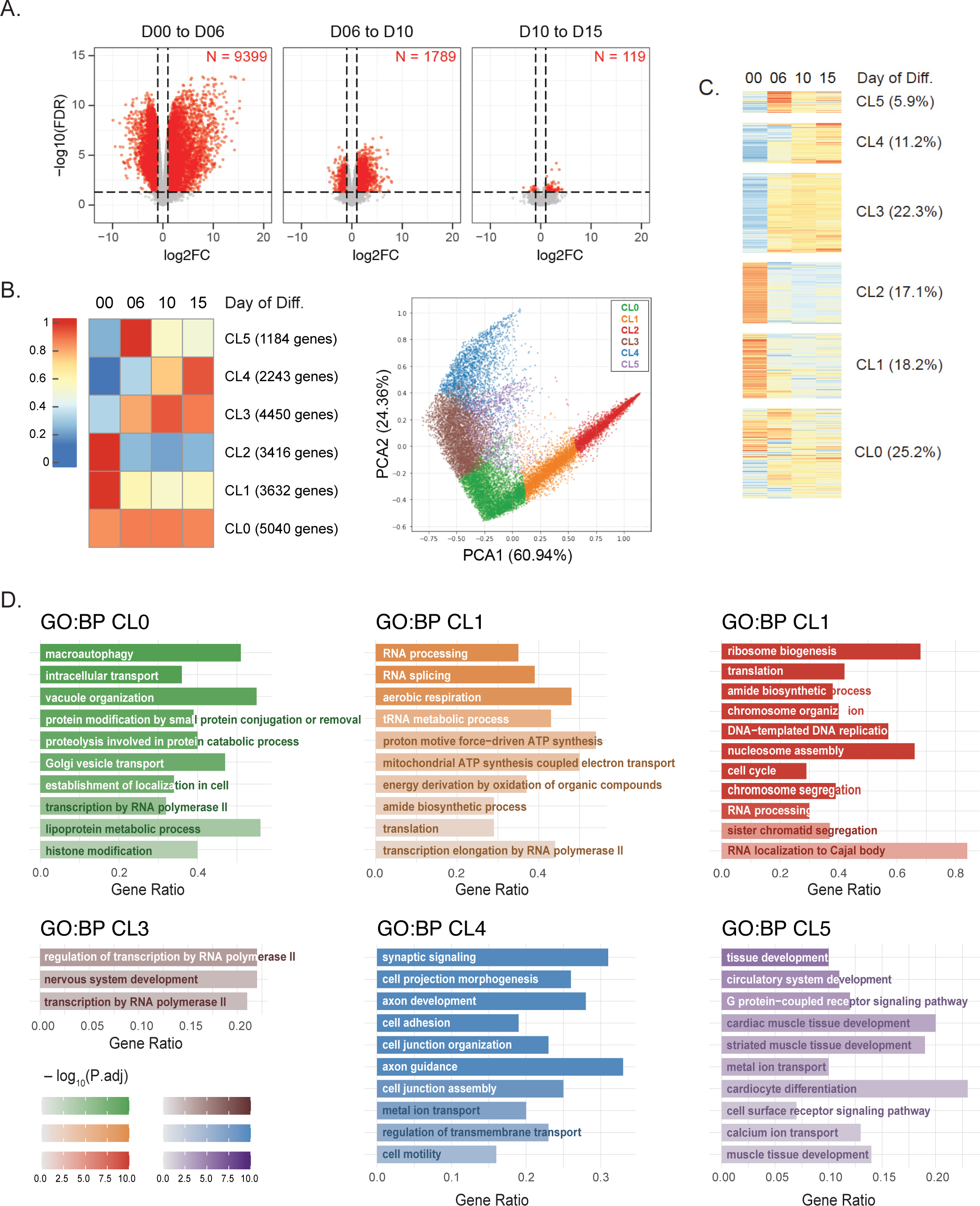
Changes in gene expression during neuronal differentiation. A: Changes in gene expression associated with the differentiation process in wild type LUHMES cells. Genes with statistically significant changes in expression between consecutive time points of differentiation are marked red. B: Clusters representing various gene expression patterns associated with the differentiation of LUHMES cells, obtained by k-means clustering of RNA-seq data. k-means of 6 resulted in good representation of expression patterns and separation of genes into clusters as seen in the PCA analysis plot on the right. Each point on the plot represents a gene, genes are colored by clusters. C: Heat map representation of expression patterns of all the genes during differentiation of WT cells, separated by clusters. D: Representation of GO terms associated with defined gene clusters, as analyzed by g::Profiler. Detailed outcome of the analysis is in the Supplementary Table 2.

**Figure 4.**
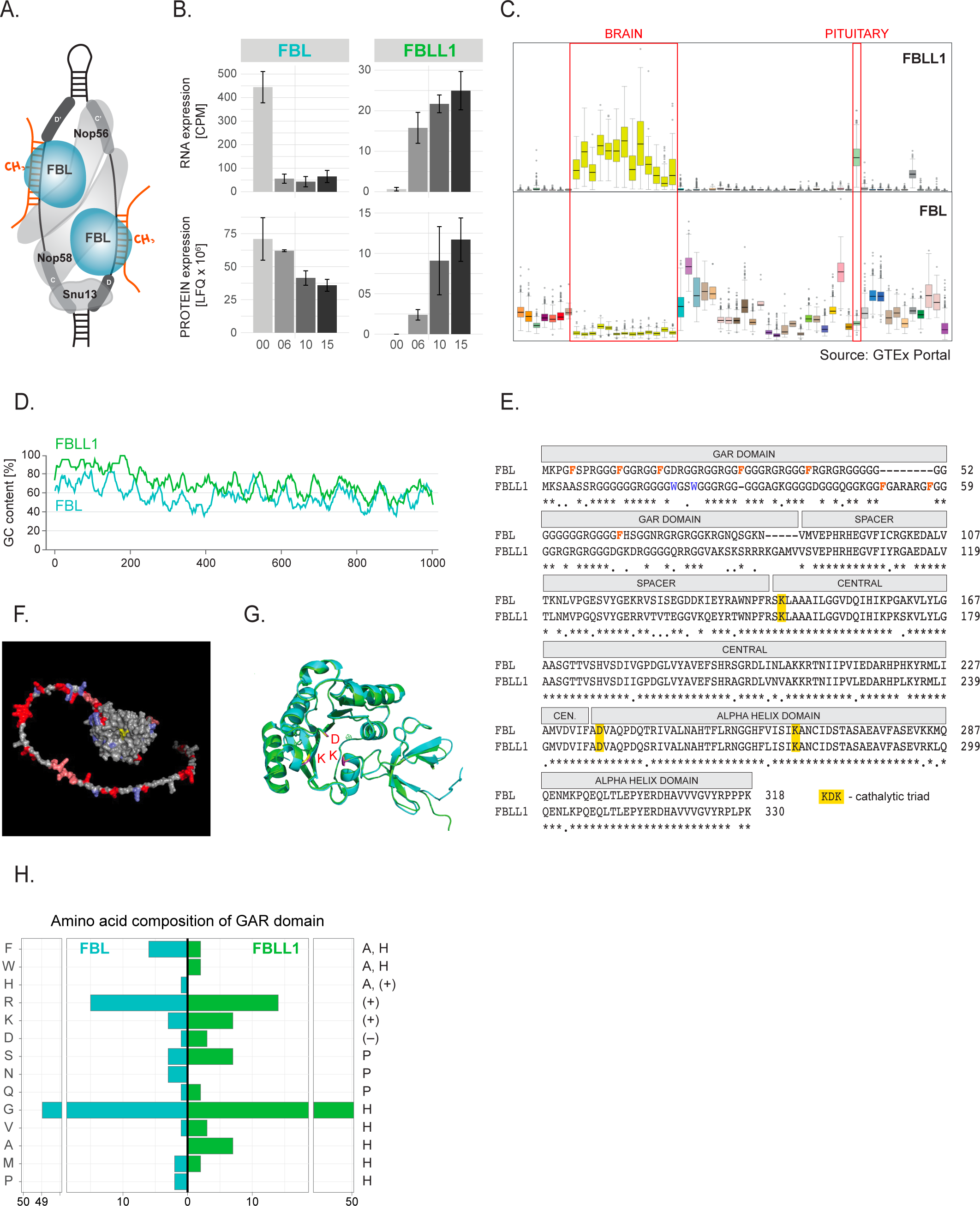
A Fibrillarin homologue is expressed during neuronal differentiation. A: A schematic structure of C/D box snoRNAs with canonical set of proteins: Fibrillarin (FBL), Nop56, Nop58 and Snu13. **B:** Changes in FBL and FBLL1 mRNA and protein abundance during neuronal differentiation. C: Relative tissue expression of FBL and FBLL1 mRNAs from GTEx portal. Red boxes mark brain tissues, including pituitary gland. D: Difference in Guanine-Cytosine (GC) content of the coding sequence of FBL and FBLL1. E: FBL and FBLL1 proteins are highly similar with the highest number of differences within GAR domain. Catalytic triad within methyltransferase active site marked yellow, is present in both proteins. Asterisk: same amino acid, dot: similar amino acid. F: Fibrillarin structure predicted by AlphaFold with indicated differences between FBL and FBLL1. Color coding: silver: the same amino acid; blue: similar amino acid, red: different amino acid; pink: deletion; yellow: catalytic triad. G: High similarity of FBL (cyan) and FBLL1 (green) proteins structures as predicted by AlphaFold and overlayed in PyMOL. Catalytic triad marked red. H: Differences in amino acid composition of FBL and FBLL1 GAR-domains. On the right of the panel, A: aromatic amino acid, H: hydrophobic, (+): positively charged, (-): negatively charged, P: polar.

To identify gene expression patterns related to neuronal differentiation we performed k-means clustering, with k=6 giving the best resolution without repeating patterns (Figs. 3B and 3C). Principal component analysis (PCA) for all quantified genes, confirmed good separation between clusters (Fig. 3B; genes are colored by cluster). The results are consistent with previous data on LUHMES cells and other analyses of neurogenesis (Lauter *et al*., 2020). The biggest cluster, CL0 (25.2 % of all genes) contains genes that are stably expressed throughout differentiation. GO term analysis using gProfiler, indicates enrichment for intercellular transport, transcription, proteolysis and macroautophagy; essential functions regardless of differentiation status (Fig. 3D; detailed results in Supplementary Table 1). CL1-3 comprise genes that substantially change expression between D00 and D06, as the LUHMES cells exit mitosis. CL1 and CL2 genes showed decreased expression (CL2 more acutely than CL1) and are involved in growth, cell cycle regulation and progression, transcription, ribosome biogenesis and translation. CL3 genes had increased expression at D06, but lack clear GO term enrichment.

The smaller CL4 (11%) and CL5 (6%) comprise genes that continued to change later during the differentiation time course. Genes from CL4 generally showed elevated expression at D06 that continued over later time points. They are enriched for characteristic neuronal functions; including neurite development, axon guidance, intercellular communication, formation of synapses, cell junctions, transmembrane transport, or cell motility. CL5 genes rose sharply at D06 and then declined. They show enrichment for tissue development and establishment of higher level organization, particularly muscles - both striated and cardiac. Decreased expression after an initial peak, potentially indicates similarities in early developmental pathways for neurons and muscles, with divergence at this point.

Notably, genes from clusters CL4 and CL5 are enriched for neuronal/developmental functions (Fig. 3D) and show regulated expression at times when SNORD115 and SNORD116 snoRNAs accumulate.

### Differentiating LUHMES cells express a homologue of core snoRNP protein Fibrillarin

Canonical box C/D snoRNAs are packaged with four proteins; the snoRNA-directed RNA 2’-*O*-methyltransferase Fibrillarin (FBL) together with NOP56, NOP58 and SNU13 (Fig. 4A). During LUMHES differentiation the mRNA and protein levels of FBL fell rapidly, probably reflecting a shut-down in ribosome synthesis as cell division stops (Fig. 4B). Expression of the other snoRNP components was also reduced, but to a lesser extent (Table S1). Unexpectedly, transcriptome analysis of mRNAs that are strongly upregulated identified an almost uncharacterized homologue of FBL designated Fibrillarin Like 1 (FBLL1) (Fig 4B). FBLL1 mRNA and protein had very low abundances in undifferentiated, cycling cells, but increased rapidly during differentiation. As expected, changes in protein levels lagged behind mRNA during FBL depletion and FBLL1 induction. Inspection of published transcriptome data (GTEx portal) strongly supported brain-specific expression of FBLL1 mRNA, which was generally anti-correlated with FBL expression (Fig. 4C; the pituitary is located within the brain). The *FBLL1* gene is unusual in lacking introns, suggesting that it may have originated via retro-transposition. It also has a notably high G+C content: 89% within the 5’ 200 nt and 71% overall (Fig. 4D). This is a feature that generally correlates with high translation efficiency (Kudla *et al*, 2006) and may offset negative effects on mRNA transport and translation due to the absence of introns from the *FBLL1* transcript.

FBL and FBLL1 share 83% similarity and 76% identity (Fig. 4E). The globular, enzymatic cores of FBL and FBLL1 are highly homologous and predicted to adopt very similar structures (Figs. 4F and 4G). Notably, residues directly implicated in RNA methyltransferase activity of FBL (KDK) are conserved in FBLL1, as are amino acids surrounding the catalytic site (Figs. 4E and 4G). Both FBL and FBLL1 share the presence of an unstructured domain, rich in glycine and arginine (termed the GAR or RGG domain) (Aris & Blobel, 1991), and the composition of this region shows differences between FBL and FBLL1 (Fig. 4E, 4F and 4H). In particular, FBLL1 has more Lys and Trp than FBL, and fewer Phe residues. These changes potentially mediate alterations in RNA binding and condensate formation (Kim & Kwon, 2021) and could be related to the predicted stabilization of SNORD116 during differentiation (see Discussion).

### RNA abundance changes in a disease model system

To understand the role of SNORD116 and SNORD115 clusters in neuronal differentiation and PWS, we precisely deleted regions of *SNHG14* gene containing the snoRNA clusters from only the paternal (expressed) chromosome, using CRISPR in LUHMES cells (Fig. S2A). The heterozygous deletion strains were designated H115 or H116, respectively. *SNHG14* is not transcribed from the maternal chromosome, which was left intact. Analysis by PCR confirmed the heterozygous deletion (Fig. S2A). Northern hybridization demonstrated the expected absence of snoRNA expression in H115 and H116 (Fig. S2B).

Most canonical box C/D snoRNAs function as modification guides for stable RNA species, predominately rRNAs, but a minority are required for correct pre-rRNA processing. Complementarity between SNORD115 nor SNORD116 and the pre-rRNA was found, but neither shows the very specific interaction pattern expected to be required for methylation (Cavaillé *et al*., 2000). The interactions of snoRNAs that promote pre-rRNA folding and processing are more heterogenous. We therefor analyzed pre-rRNA processing in the wildtype, H115 and H116 strains by northern hybridization (Fig. S3). Comparison of D00 with D10 showed some reduction in pre-rRNA abundance during differentiation, consistent with the exit from cell division. However, no clear differences were observed between WT, H115 and H116 cell lines.

To assess the effects of the deletion of SNORD115 and SNORD116 clusters on gene expression, we performed RNA sequencing as above at D00 to D15. RNA-seq analysis confirmed the accurate deletion of the entire SNORD116 region from H116 and the SNORD115 region from H115 (Fig. 5A). No other clear changes in *SNHG14* gene expression were seen in the region surrounding the SNORD116/115 clusters. The total number of reads mapping to *SNHG14* was reduced in H116, consistent with the deleted region (Fig. 5C), indicating that transcription *per se* was not affected. Expression of *UBE3*, which is adjacent to *SNHG14* but transcribed from the opposite strand, was also unaffected (Fig. 5C). In the wider PWS locus region, alterations in expression were observed (Fig. 5B). *MAGEL2* mRNA was under-accumulated at D10 and D15 in both deletion strains, with a greater effect in H116 (Fig. 5D). The *MAGEL2* gene, which is causal for Schaaf-Yang syndrome, is transcribed on the opposite strand from *SNHG14* and located around 1.5 Mb upstream. Other mRNAs transcribed from the PWS region were also altered in both mutant cell lines; *OCA2* showed reduced and *GABRA5* increased expression, relative to the wild-type (Fig. 5D).

**Figure 5.**
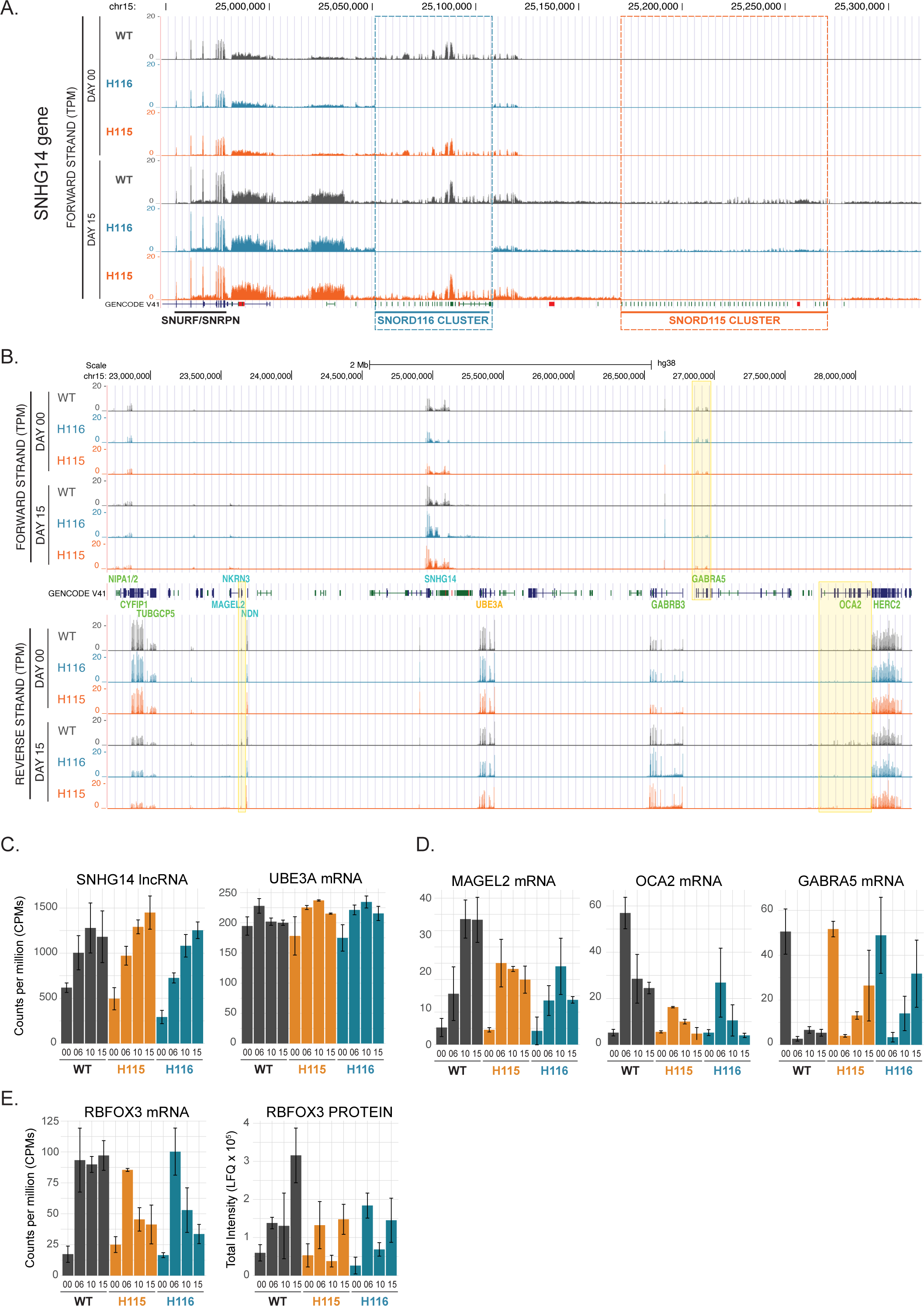
Local effects of SNORD115 and SNORD116 cluster deletions. A: UCSC genome browser view of transcription across SNHG14 gene in wild-type and mutant cell lines at D00 and D15 with indicated deletion regions. B: UCSC genome browser view of transcription across PWS locus in wild-type and mutant cells. Marked are the genes with altered expression in the deletion mutants. GABRA5 and OCA2, but not MAGEL2 are passing statistical significance thresholds. C: Expression of SNHG14 lncRNA and UBE3A, convergent gene overlapping with SNHG14, during differentiation in wild-type and mutant cells. D. mRNA expression from MAGEL2, GABRA5 and OCA2 genes from PWS locus is affected by the deletion of SNORD115 and SNORD116 clusters. E: Expression of mRNA and protein from RBFOX3 gene, with possible role in splicing of SNORD115 and SNORD116 snoRNAs from SNHG14 lncRNA introns.

Also of note, expression of RBFOX3 (NeuN) mRNA was strongly increased in the wildtype between D00 and D06, and then remained high, whereas its abundance notably declined at D10 and D15 in both H116 and H115 strains (Fig. 5E). RBFOX3 protein accumulation was delayed relative to the mRNA, as expected, but was clearly reduced in the mutant cells. RBFOX3 was previously implicated in snoRNA excision from mouse transgenes (Coulson *et al*., 2018).

Transcriptome-wide differential expression analysis revealed that for the majority of mRNAs changes during differentiation were similar in the wildtype, H115 and H116 (Fig. 6A). However, across all stages, 731 transcripts were significantly altered between wildtype and H116 cells, with 411 transcripts altered between wildtype and H115; of these 295 were common (Fig. 6B; Supplementary Table 3). Hierarchical clustering (Fig. 6C) confirmed that at D00 and D06 the wild type and mutant lines cluster together, although the mutants are already more similar to each other. At D10 and D15 with mutants cluster together, away from the wild type.

**Figure 6.**
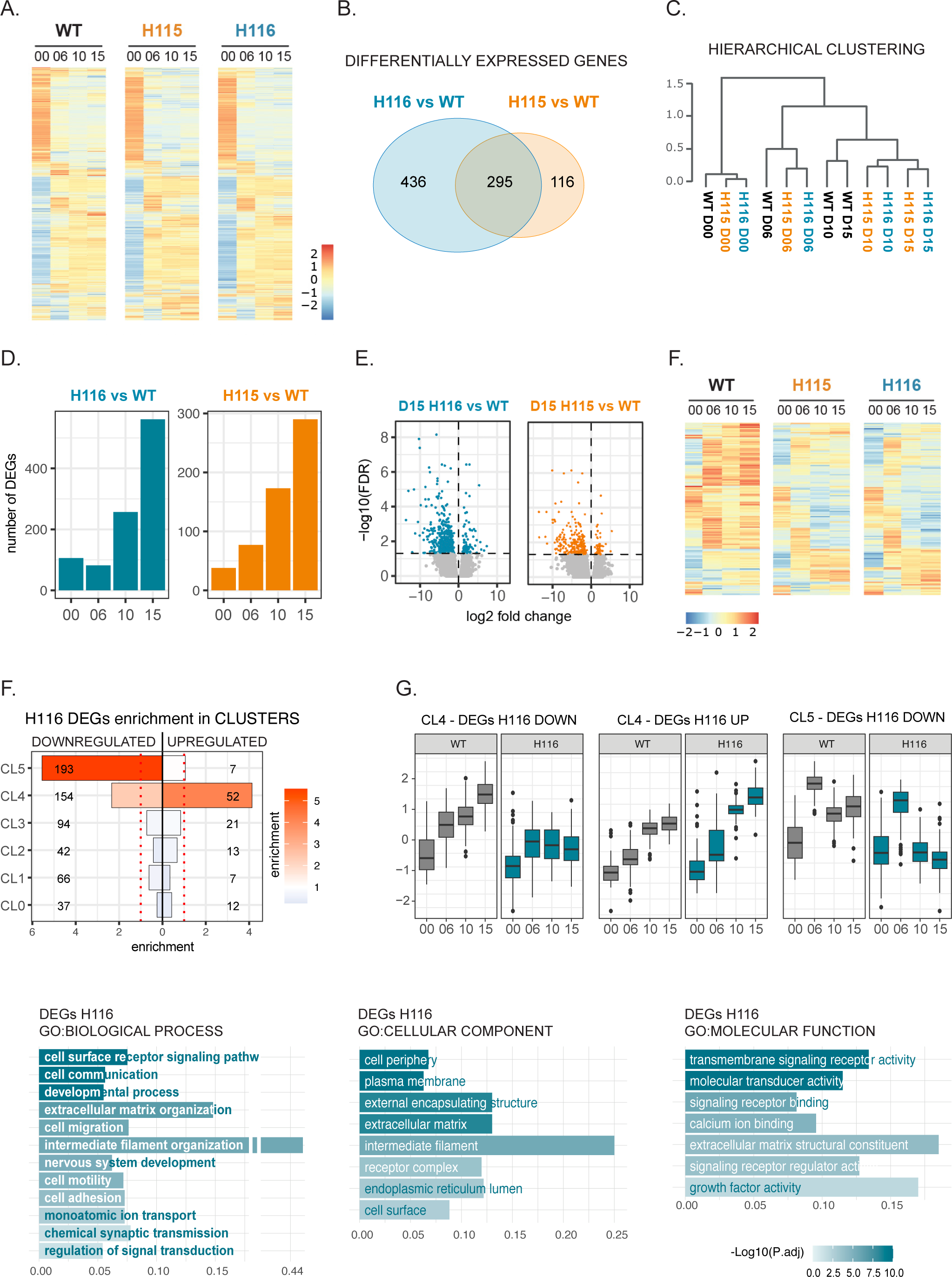
Effects of SNORD115 and SNORD116 cluster deletions on transcriptome. A: Heatmaps representing global changes to the transcriptome in differentiating wild type and mutant cell lines. Average expression (CPM) values are scaled for each gene across all the timepoints and all cell lines. B: Hierarchical clustering of RNA seq samples shows high similarity between undifferentiated cells and subsequent separation between WT and deletion mutant cell lines. C. Changes to gene expression in mutant vs wild type cells at day 15 of differentiation, indicating tendency towards decreased expression. D: Gene expression profiles of differentially expressed genes (DEGs), show very high similarity between H115 and H116 cell lines. Heat map covers all 847 DEPs identified independently in both mutant cell lines and at all timepoints. E: Number of differentially expressed genes between wild type and mutant cells increases with the progress of differentiation. F: Overlap between DEGs for H115 and H116 cell lines. G: Enrichment of DEGs in H116 cell line, in 6 clusters defined by k-means clustering (Fig. 3), based on the expression patterns of genes in wild type cells. H: Pattern of expression of H116 DEGs in wild type and mutant cells in, most affected by deletion, clusters 4 and 5. I: A summary of the GO term enrichment analysis for H116 DEGs from g::Profiler. Full outcome of the analysis is in the Supplementary Table 4.

Consistent with the hierarchical clustering, the number of RNAs differentially accumulated between mutant and WT cells lines was highest at D15 (Fig. 6D). This was notably later than most changes related to differentiation (Fig. 3A). Altered expression was seen for both mRNAs and lncRNAs, with a predominance of reduced expression (Figs. 6E). Intriguingly, some changes in mRNA accumulation were detected in undifferentiated cells lacking SNORD115 or SNORD116 clusters (106 in H116 and 38 in H115, 21 common). This was unexpected, especially for H115 cell line, as undifferentiated LUHMES cells apparently lack transcription across the SNORD115 cluster. The basis of this is unclear but could reflect the effects of deletions in the *SNHG14* gene or long ncRNA transcripts.

Notably, there was considerable overlap between transcripts with altered expression in the H115 and H116 cell lines (35% of all differentially expressed genes; DEGs) (Figs. 6B). Direct comparison of H115 and H116 revealed only 30 RNAs with significant differential expression at D15 (18 down and 12 up). These similarities were also visible when comparing heat maps for differentially expressed genes in H116 and H115 (Fig. 6E). We conclude that SNORD115 and SNORD116 are likely to functionally interact. In view of these partially overlapping phenotypes, some subsequent analyses focused only on the H116 cell line.

### Clustering of differentially expressed genes

The 6 clusters defined from RNA expression during WT differentiation (Fig. 3B and 3C) were compared to RNAs showing altered accumulation in H116 (Fig. 6F). DEGs in H116 were enriched in the two smallest clusters CL4 (30% of DEGs) and CL5 (29% of DEGs), which largely comprise neuronal related genes regulated at later stages of differentiation. Downregulated genes were enriched in both CL4 and CL5, whereas upregulated genes were clearly enriched only in CL4. Including time course data (Fig. 6G) showed that RNA levels in mutant and wild type cells are similar between D00 and D06, but diverge through D10 and D15. For the DEGs, GO term enrichment indicated processes characteristic of developing neuronal cells: regulation of membrane potential, axonogenesis, response to cAMP, endocrine system development; with highest enrichment for “extracellular matrix organization”. Enriched terms in “Cellular Compartment” indicated association with membranes, secretion and cell-cell junctions.

Particularly clear enrichment was seen for changes in mRNAs encoding intermediate filament components of the cytoskeleton / axoskeleton, which is greatly remodeled during the structural changes required for neuronal development [reviewed in (Bott & Winckler, 2020)]. Several, but not all, mRNAs encoding neurofilament components were increased during differentiation in the mutants (see Fig. S4A and legend). These included NEFL (Neurofilament Light Chain), which is the most abundant neurofilament component. NEFL is linked to disease and is used as a biomarker for neuronal damage. In marked contrast, expression of almost all of the large family of keratins and associated proteins was strongly reduced in the mutants (Fig. S4B).

We conclude that RNAs showing altered expression in the deletion mutants were generally subject to regulated expression later in wild-type neuronal development. This is consistent with the time-course of snoRNA accumulation.

### *SNORD* deletion does not clearly alter pre-mRNA splicing

Previous reports proposed roles for SNORD115, SNORD116 and extended snoRNA-related ncRNAs in alternative pre-mRNA splicing, acting directly via base-pairing with the target pre-mRNA or through protein sequestration (Baldini *et al*., 2022; Bazeley *et al*, 2008; Bochukova *et al*., 2018; Kishore & Stamm, 2006; Yin *et al*., 2012). We therefore analyzed our RNAseq data for changes in pre-mRNA splicing using DEXSeq (Anders *et al*, 2012). During differentiation in wildtype cells (comparing D00 with D15) we identified multiple alternatively spliced transcripts, showing this phenomenon to be frequent (for an example see Fig. S5A). We next compared the deletion cell lines with the wildtype at D15. This identified a relatively small number of potentially alternatively spliced introns (95 for H116 and 73 for H115). The corresponding RNAseq data for each was inspected visually in the UCSC genome browser. Surprisingly, only 9 genes showed clearly altered expression of a subset of exons in H116 and H115 cells, all with quite small effects. Moreover, for all of these genes differential expression most likely reflected the use of alternative transcription start sites (*OLFM1, MYO15A, NRXN1, IQSEC1, NAV1* and *GSE1*) or alternative termination sites (*LAMP2, GNAO1* and *HERC2P3* pseudogene - a frequent breakpoint in PWS patients) rather than the alternative splicing *per se* (Fig. S5B). We note that all candidate genes were affected very similarly in H116 and H115 mutant cell lines, making it less likely that sequence-specific targeting of pre-mRNAs by these snoRNAs directs alternative splicing. We also visually inspected the RNAseq data for multiple other genes previously predicted or reported to be regulated by SNORD116; no changes were confirmed in our data. The *HTR2C* gene (Kishore & Stamm, 2006) was not detectably expressed in LUHMES cells. We conclude that alternative splicing is unlikely to underlie the changes in gene expression in the *SNORD115* or *SNORD116* deletion cell lines.

### Changes in translation in *SNORD* deletion lines

SNORD115 and SNORD116 derived ncRNAs might also influence mRNA translation; potentially via post-transcriptional effects on mRNP composition and/or nuclear cytoplasmic transport. To assess this, the total proteome was determined for the wildtype, H115 and H116 cell at D00, D06, D10 and D15 using HPLC-coupled, tandem mass-spectrometry with data-independent acquisition (DIA) (Fig. S6A, and S6B, Supplementary table 1 and 5).

Proteomic data broadly replicated the results from transcriptomic analyses: (1) Large and rapid changes during the first interval of differentiation (Fig. S6A and S6B). (2) High similarity between undifferentiated wildtype and mutant cells, with progressive divergence during the differentiation process (Fig. S6A). (3) Increased numbers of differentially expressed proteins (DEPs) between mutant and WT cells over the time course of differentiation (Fig.S6D). (4) Substantial overlap between proteins with altered abundance in H115 and H116 cells (Fig. S6C). Notably, in hierarchical clustering the corresponding proteomic intensities and transcriptomic read counts were grouped together, supporting their accuracy (Fig. S6E).

Proteomic and transcriptomic data were compared to identify proteins showing increased or decreased abundance relative to the corresponding mRNA in mutant cells. As expected, the relationship between protein steady state level and RNA steady state level (P/R ratio) is highly variable for different genes: The highest P/R ratios were for TUBB4A (3 x 10^9^), ACTA1 (9 x 10^8^) and MAP1LC3A (6 x 10^8^), with the lowest for MT-CO1 and MT-CO3 subunits (56 and 93, respectively) (Fig. S6G). Spearman’s correlation between protein and mRNA expression levels in steady population of undifferentiated cells ρ = 0.55 and in differentiating cells ρ < 0.4 (Fig. S6F).

For individual genes P/R ratios were surprisingly stable, especially when compared within the same stage of differentiation. For most genes P/R fold difference between mutant and wildtype cells oscillated closely around 1 (Fig. S1H). Spearman’s correlation between P/R ratios was also very high, ρ > 0.9 within the same stage of differentiation (Fig. S6I) and ρ > 0.8 between different stages (data not shown). We therefore used P/R ratio to identify potential instances of different post-transcriptional gene regulation in wildtype and mutant cells. We focused on DEPs with the most statistically significant changes in protein expression and at least 2-fold difference in P/R ratio in at least two differentiation stages, for either SNORD cluster deletion (Fig. S6I). In total 90 genes passed these thresholds; 37 with increased P/R ratio and 53 decreased. A summary of RNA and protein expression, together with fold change differences in expression between wildtype and mutant cells is presented as a heatmap (Fig. 7A). More detailed data, with error bars, are shown for selected genes in Fig. 7B. Several genes showed stable mRNA levels but pronounced differences in protein abundance.

**Figure 7.**
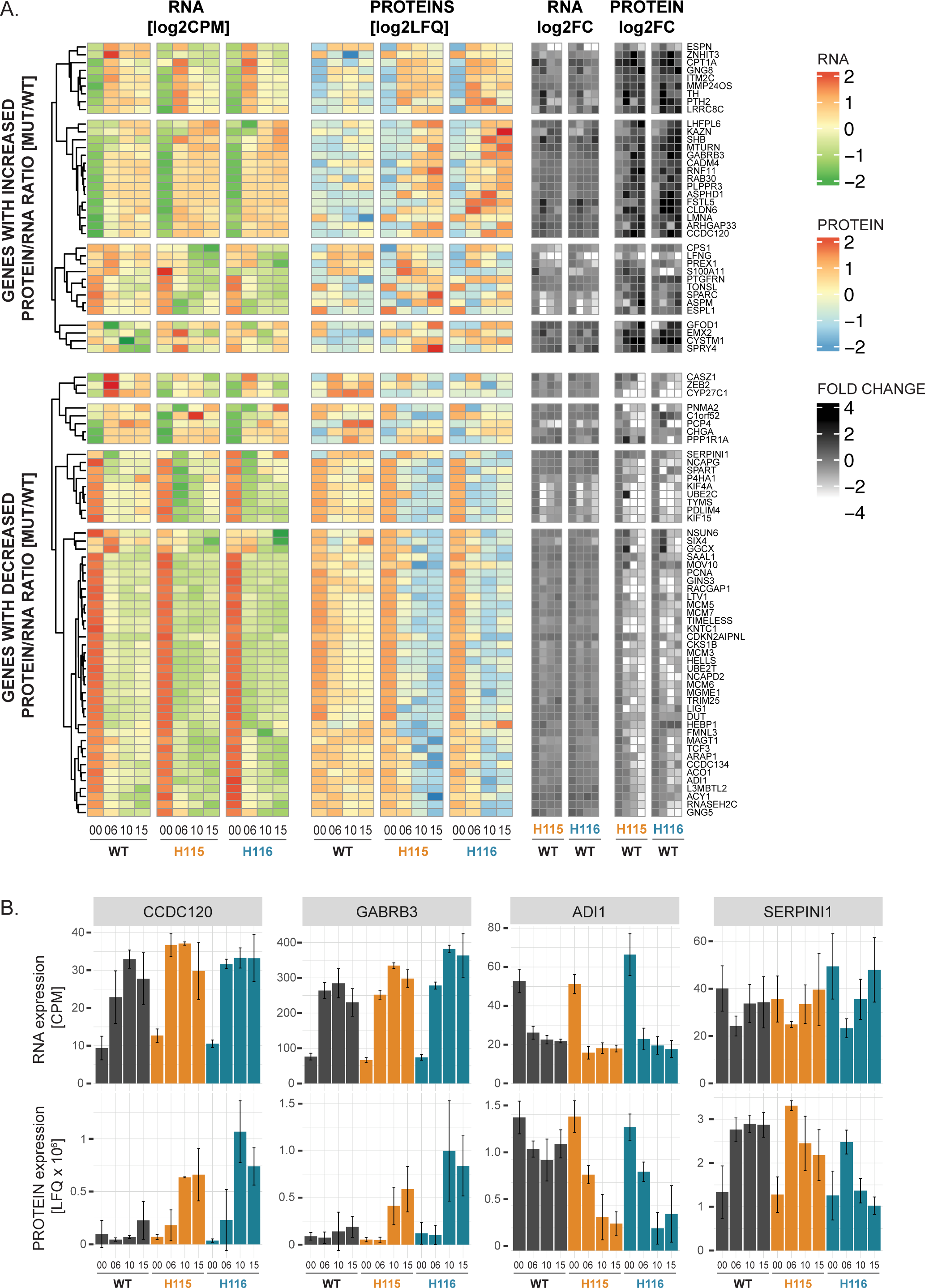
Post-transcriptional effects of SNORD115 and SNORD116 clusters loss. A: Combined analysis of transcriptome and proteome data revealed different relation between protein and mRNA expression in wild type vs mutant cells. Protein/RNA (P/R) ratio can be both decreased and increased, and can be a sign of the involvement of SNORD115 and SNORD116 clusters in post-transcriptional regulation of gene expression. B: mRNA and protein expression plots for selected genes displaying altered P/R ratio in wild type and mutant cells.

We conclude that deletions of *SNORD115* or *SNORD116* alter protein abundance relative to mRNA levels, with considerable overlap in targets. Direct effects of ncRNAs on cytoplasmic protein stability appear unlikely. We therefore infer, direct or indirect, differences in mRNA translation efficiencies.

### Altered developmental timing in the absence of the SNORD116 cluster

The above data all indicate that undifferentiated WT and mutant cell lines are very similar but diverge over the course of differentiation, particularly through D10 and D15. To better characterize the changes in gene expression that underlie separate clustering of WT, H115 and H116 samples, the transcriptome data was subjected to principal component analysis (Fig. 8A) (Pearson, 1901). At D00, the undifferentiated wild-type, H115 and H116 cells all cluster together. By D06 some separation is observed between wild-type and mutants, which becomes progressively more marked at D10 and D15. By inspection, it appeared possible that the wild-type and mutant cells were on a similar trajectory, but with the mutants advancing further. Analysis of proteome data also gave results consistent with this hypothesis.

**Figure 8.**
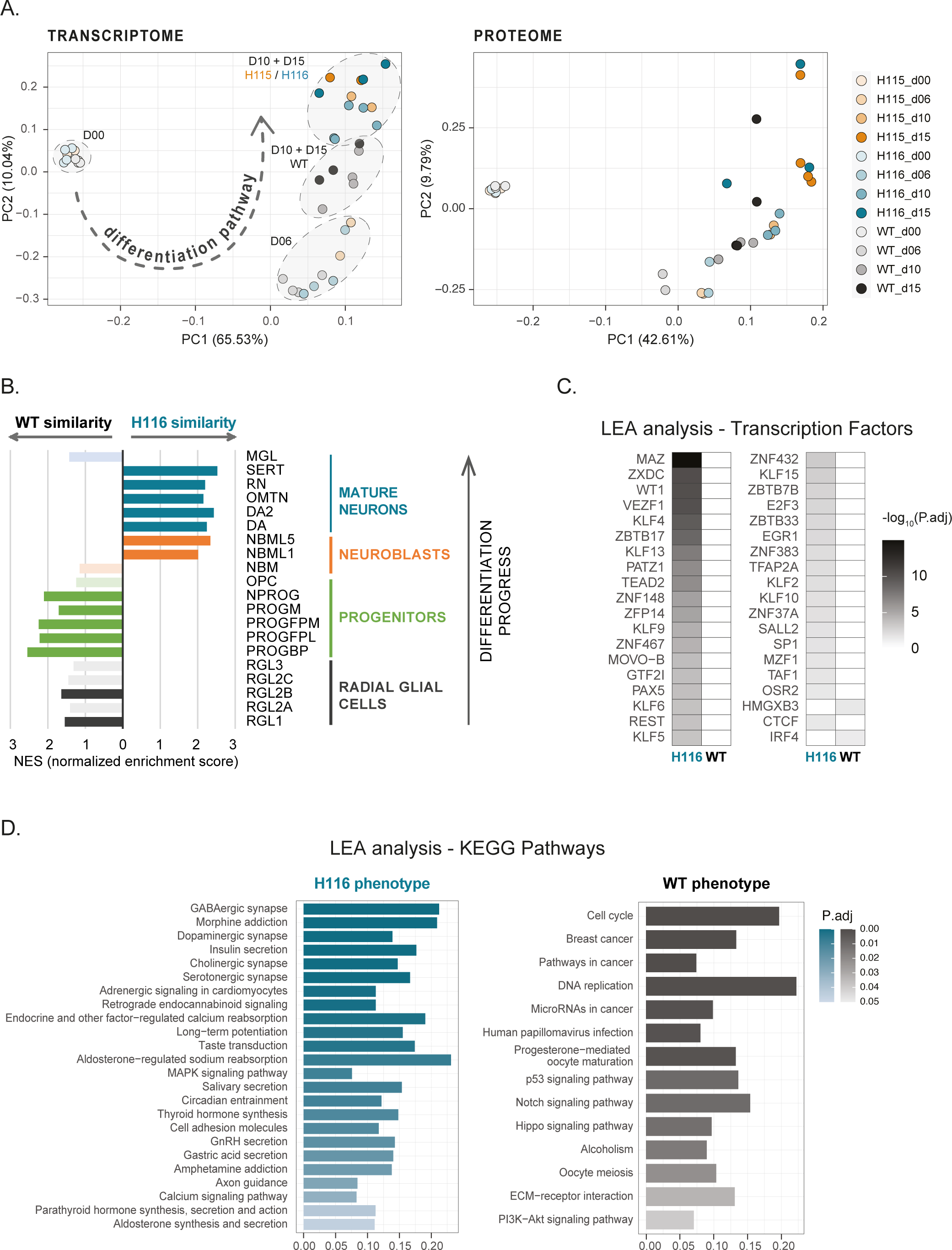
Neurons lacking SNORD116 show accelerated differentiation. **A**: PCA analysis of RNA-seq data implies accelerated differentiation of H115 and H116 cells lines along the assumptive differentiation pathway. This trend is supported by the PCA analysis of proteomic data. **B**: Comparison of H116 and WT transcriptomes at D15 of differentiation with transcriptomes of various cell types identified during human midbrain development (scRNA-seq) by Manno et al., suggests that mutant cells exhibit more “mature” phenotype than wild-type cells; GSEA analysis. RGL1-3: radial glia-like cells; PROG: progenitor cells, BP: basal plate, FPL: lateral floorplate, FPM: medial floorplate, M: midline; NPROG: neuronal progenitors; OPC: oligodendrocyte precursor cells; NBML1-5: mediolateral neuroblasts; DA-DA2: dopaminergic neurons; OMTN: oculomotor and trochlear nucleus; RN: red nucleus; SERT: serotonergic; MGL: microglia. **C**: Expression of genes responsible for “mature” phenotype of H116 cells is associated with the activity of transcription factors with high significance. There is no similar association with the genes responsible for wild type phenotype. Genes associated with phenotypes were identified by GSEA/LEA analysis. Transcription factor enrichment analyzed by g::Profiler. Detailed outcome of the analysis is in the Supplementary Table 6. **D**: KEGG pathways enrichment analysis of the LEA identified genes, responsible for separation between H116 and wild-type phenotypes, as analyzed by g::Profiler. Detailed outcome of the analysis is in the Supplementary Table 6.

To explore this hypothesis, gene expression at D15 of differentiation in wildtype and H116 cells was compared to a large set of mid-brain derived neuronal cell types. Gene Set Enrichment Analysis (GSEA) (Subramanian *et al*, 2005) was applied using published data (La Manno *et al*, 2016) as deposited in the Molecular Signatures Database (MSigDB) (Subramanian *et al*., 2005). In Figure 8B, cell types identified during embryonal development of the human midbrain have been ordered with the least mature and the foot and the most mature at the top. Strikingly, the wild-type LUHMES cells more closely correlated with the more immature neurons, whereas the H116 cells showed greater similarity to more mature cell types. Representative enrichment plots for the analysis are shown in Figure S6.

The subset of genes that contributed most to these distinctions were identified by leading-edge analysis (LEA; Supplementary Table 6). We aimed to determine whether they are *prima facie* candidates to be potentially disease linked. Associated KEGG pathways were identified using gProfiler (Fig. 8C) The WT phenotype was associated with enrichment for the cell cycle, DNA replication and various cancers (presumably reflecting growth-related activities). In contrast, the H116 phenotype was associated with neuronal activity: synapses, calcium signaling and hippocampal long-term potentiation. We note that several terms enriched for H116 correlate with PWS phenotypes; including hormonal regulation [(GnRH (Miller *et al*, 2009), parathyroid hormone (Iughetti *et al*, 2019), aldosterone (Kusz & Gawlik, 2022)], insulin secretion (Lautala *et al*, 1986), salivary secretion (Hart, 1998), circadian entrainment (Powell *et al*., 2013) and addictive behavior (Salles *et al*, 2021).

A surprisingly large number of genes contribute to separating the “more mature” H116-phenotype from the “less mature” WT-phenotype. We therefore considered the possibility that transcripts from SNORD116 cluster affect availability of transcription factors (TFs) that regulate expression of multiple target genes. We performed g:Profiler multiquery analyses against transcription factors from the TRANSFAC database (Wingender *et al*, 2000) on genes from the LEA; including 493 genes contributing to H116 phenotypes and 503 genes contributing to WT phenotypes.

Notably, genes linked to the wildtype were associated with few TFs, whereas H116-linked genes were associated with many TFs (Fig. 8D; Supplementary Table 6). Relevant TFs mostly showed similar levels for mRNA and protein (where detected) between WT and mutants. Exceptions were: TEAD2/ETF (reduced at D10 and D15), PAX5 (reduced at D06-D15, particularly in H115) and KLF6/CPBP (increased at D06-D15). These observations would be consistent with increased activity for several TFs in the absence of SNORD cluster derived ncRNAs.

We conclude that the data support a model that the loss of SNORD116 advances developmental progression. The mRNAs specifically affected show enrichment for neuronal function, including phenotypes related to features of PWS.

## DISCUSSION

### Changes in RNA and protein levels during neuronal development

We initially followed the pattern of transcription from the ∼600Kb, PWS-associated, *SNHG14* locus during neuronal differentiation. Histone modification data (see Fig. 2B) indicates that the entire *SNHG14* region forms a single transcription unit. However, we clearly distinguished regions showing differential transcript accumulation: SNURF/SNRPN region, SPA1 region, SNORD116 cluster and SNORD115 cluster, the latter two being our major focus. During differentiation, transcriptome data revealed substantial accumulation of spliced *SNHG14* exons relative to introns, and little change in exon abundance within the SNORD116 region. This indicated that transcription and splicing of the *SNHG14* primary transcript is essentially unaltered during differentiation. We detected previously reported sno-lncRNAs, long ncRNAs containing two SNORD116 snoRNAs joined by a linker (Sledziowska *et al*., 2023; Wu *et al*., 2016; Yin *et al*., 2012). During differentiation, accumulation of sno-lncRNA 1-3 decreased, sno-lncRNA 4 was unchanged, while sno-lncRNA 5 was not detected. In contrast, mature SNORD116 snoRNAs were strongly accumulated, presumably reflecting regulated stabilization after excision from spliced introns. This expression pattern would be consistent with competing maturation pathways for SNORD116 snoRNAs and sno-lncRNAs.

A quite different pattern was seen for the SNORD115 region, located further 3’. Very low levels of all transcripts were seen in undifferentiated cells, with accumulation of both lncRNA exons and mature snoRNAs only during differentiation. We cannot exclude the possibility that the primary transcript encompassing this region is rapidly and completely degraded in undifferentiated cells. However, it seems more likely that transcription terminates immediately prior to the SNORD115 region. During differentiation, termination readthrough increases with immediate splicing and SNORD115 accumulation.

We speculate that accumulation of mature SNORD115 and SNORD116 snoRNAs is related to the differentiation driven appearance of FBLL1 protein. FBLL1 is a close homolog of FBL, a core component of canonical snoRNPs, but is uncharacterized and apparently neuron-specific. The main differences lie within the GAR/RGG domain, a low complexity, glycine plus arginine rich, RNA interacting region, also present in several other nucleolar proteins. The GAR domain of FBLL1 has fewer charged arginine residues than FBL, but more lysine, and fewer aromatic, phenylalanine residues, but more tryptophan. The uncharged, planar amino acids can intercalate into or stack onto RNA stems. RNA interactions by the GAR domains are precited to be largely sequence independent but are likely to facilitate condensate formation (Kim & Kwon, 2021). The changes between FBL and FBLL1 may impart different phase separation properties to snoRNP complexes. We note that key steps in snoRNP maturation take place in phase-separated Cajal bodies (Verheggen *et al*, 2002), offering a potential link. Functional analyses of FBLL1 will be reported elsewhere.

To better understand PWS-related changes, we created two heterozygous deletion cell lines lacking expression from either the SNORD116 or SNORD115 cluster. Notably, the LUHMES cells used have “never” expressed these snoRNAs (or at least have not done so for many generations), but then express the ncRNAs from the endogenous locus, in an appropriate cell type, during differentiation. We therefore expected to better identify primary effects, before compensatory mechanisms become dominant.

Upon deletion of the SNORD116 or SNORD115 clusters, accumulation of neighboring exons was unaffected, with no indication of unusual transcripts generated from *SNGH14*. Within the PWS region, genes in proximity to *SNHG14* gene were also unaffected. We concluded that deletion of SNORD115 and SNORD116 clusters did not influence transcriptional regulation within in PWS locus. There were, however, changes the wider PWS locus. Notable amongst these was the under-accumulation of mRNA from *MAGEL2,* located ∼1.5 Mb “upstream” from *SNORD116*. Mutations in *MAGEL2* are causal in Schaaf-Yang syndrome, which shows striking similarities to the PWS-phenotype (Chen *et al*, 2020; Runte *et al*., 2001). We predict that reduced *MAGEL2* mRNA and protein will contribute to PWS phenotypes in individuals with *SNORD116* microdeletions. However, we did not detect MAGEL2 protein in our proteomic data, suggesting that its depletion does not cause cellular defects reported here.

Throughout the transcriptome numerous genes and ncRNAs showed altered abundance in the mutant cell lines. In the wild type, changes in mRNA abundance during differentiation predominately occur during the initial period (to D06) as the cells exit mitosis. In contrast, differences between the wild type and mutants predominately occurred at later stages, consistent with the time course of snoRNA accumulation. We noted a striking degree of overlap between RNAs with altered abundance in the SNORD116 and SNORD115 deletion strains. Although SNORD115 and SNORD116 are both classed as box C/D snoRNAs, they lack clear homology outside the conserved structural features of the C/D and C’/D’ elements. They would not therefore be expected to base pair to the same sequences. We speculate that the SNORD115 and SNORD116 families may share overlapping, but non-redundant target RNAs, or functionally interact in some way – perhaps acting cooperatively on some targets. A previous report found potential interactions between SNORD115 and SNORD116 when ectopically expressed in non-neuronal, human HEK cells (Falaleeva *et al*., 2015).

Mature SNORD115 and SNORD116, as well as extended snoRNA transcripts, were previously reported or predicted to affect alternative pre-mRNA splicing (Baldini *et al*., 2022; Kishore & Stamm, 2006; Yin *et al*., 2012). To pursue this, RNA sequencing was performed at considerable depth and cases of differential intron accumulation were identified. These were, however, few in number and the clearest changes appeared to actually reflect altered transcription initiation or termination events. We were unable to reproduce any previously described splicing changes in our system. Accumulation of the neuronal-specific splicing factor RBFOX3 (NeuN), which participates in splicing of introns containing SNORD116 in mouse transgenes (Coulson *et al*., 2018) was reduced two-fold at the level of RNA and protein. This was not associated with a clear splicing defect, but the level of expression in LUHMES might be too low high for an evident phenotype.

In marked contrast, many examples of apparent alterations in translation efficiency were discovered by comparison of transcriptome and proteome data in the wild type and mutant lines during differentiation. This identified numerous proteins with increased or decreased abundance relative to the corresponding mRNA levels. We speculate that translation efficiency is altered by changes in mRNP composition and/or nuclear/cytoplasmic transport following ncRNA loss. There was substantial overlap in the effects of deletion of the SNORD116 or SNORD115 clusters, suggesting overlapping functions.

### Altered gene expression in cells lacking SNORD116 or SNORD115

Induction of differentiation very quickly induces huge changes to the transcriptome. In wildtype LUHMES cells, about 50% of the quantified transcripts increased or decreased their expression between D00 and D06, as the cells left mitosis and entered into post-mitotic differentiation. However, clustering analyses identified two relatively small groups of transcripts (CL4 and CL5) that showed regulated changes also later in differentiation.

Comparing the wildtype and mutant H116 and H115 cell lines, we noticed that the undifferentiated cells are very similar, but they progressively diverge with differentiation. Intriguingly, most of the differentially expressed genes disclose late in our time course (D15) when expression of the most genes has already stabilized. Differentially expressed genes mainly fall into the CL4 and CL5 clusters, underlining non-random selection of ncRNA sensitive genes.

Differentially expressed genes were associated with molecular functions closely related to neuronal differentiation and functioning, such as axon guidance, regulation of membrane potential or response to cAMP. Interestingly, they were also highly associated with cell surface and extracellular matrix (GO term: cellular component) and 140 of 350 mRNAs downregulated at D15 in H116 cells are annotated as glycoproteins (https://www.uniprot.org/keywords/KW-0325). A recent report described the detection of glycosylated RNAs, including SNORD116 (Flynn *et al*, 2021). The significance of this observation remains unclear, but it suggests a possible link between SNORD116 and glycosylation pathways.

Particularly striking enrichment was identified for mRNAs encoding components of intermediate filaments, which must be substantially remodeled during the large-scale changes in cell morphology required for neuronal development. Almost all keratins and keratin-associated mRNAs were strongly depleted (Fig. S4A and B), whereas neurofilament mRNAs were generally upregulated (Fig. S4B). This may be linked to the changes in developmental timing described below.

### Altered developmental progression in cells lacking SNORD116

Principal component analysis of the RNA and protein expression data suggested that the mutant and wild-type cell lines might be on the same trajectory, but with the mutants advancing more rapidly. Comparison of mRNA levels in wild type and H116 line to a range of mid-brain cell types provided strong support for this interpretation; cells lacking SNORD116 showed greater similarity to more mature neurons, while the wild type resembles more immature cell types.

Moreover, mRNAs that are specifically affected show enrichment for neuronal function, including phenotypes related to features of PWS. It might be envisaged that many different defects in cellular metabolism could slow development, perhaps by limiting gene expression. In contrast, advances in developmental progression seem likely to reflect more specific changes.

Many genes contributed to the “more mature” phenotype of snoRD116-depleted neurons, and multiple TFs were predicted to participate in their regulation. Among these, we note that the MYC-associated zinc finger protein (MAZ) is activated by RNA via EWS RNA-binding protein 1 (EWSR1) (Hoell *et al*, 2011; Li *et al*, 2019), suggesting the possibility of regulation by the snoRNAs. Overall, the data indicate that the ncRNAs may act to reduce TF activity.

The effects on gene expression reported here are much more marked than in several previous analyses. This might reflect the fact that LUMES cells natively express SNORD115 and SNORD116 during differentiation. They may therefore be “primed” for the appearance of these snoRNAs, expressing the corresponding cofactors and targets. In addition, like cells in early development, LUHMES cells have never previously expressed, or required, the snoRNAs, thus avoiding selection for compensatory pathways. Detection of the changes in gene expression timing reported here were greatly facilitated by the rapid and highly synchronous development of LUHMES cells, but would potentially be obscured by heterogeneity in other model systems.

We speculate that, in the absence of timely, neuronal SNORD116 expression, early developmental timing is altered. This may generate neuronal sub-populations that are out of synchrony in individuals lacking SNORD116. Resulting impairments in optimal network generation, may trigger functional consequences later in development. However, by this time the original primary problem will not be readily identified. Previous mouse models for PWS have not fully recapitulated the human disease phenotype. We note that the timing of mouse brain development is substantially different from humans, possibly explaining these difficulties.

Developmental timing is crucial for brain development; key processes including cell division, migration and the establishment of cell-cell contacts are separated in time and depend on the extracellular environment. In humans brain development is slow relative to growth, compared to many other animals, allowing large brains, with maturation continuing even after birth. It is well documented that individuals with PWS have significant structural brain alterations. We postulate that these may fundamentally reflect subtle changes in the progression of neuronal maturation in early development.

## MATERIALS AND METHODS

### Cell culture

LUHMES cells (ATCC cat# CRL-2927) (kindly supplied by A. Bird) were cultured according to published protocols (Scholz *et al*, 2011; Shah *et al*., 2016)). Briefly, cells were grown on poly-L-ornithine (PLO) and fibronectin precoated dishes; for proliferation in Advanced DMEM/F12 (ThermoFisher Scientific, 12634028) with addition of L-Glutamine, N2 (ThermoFisher Scientific, 17502048) and βFGF (R&D Systems, 4114-TC-01M); for differentiation in Advanced DMEM/F12 with addition of L-Glutamine, N2, GDNF (R&D Systems, 212-GD-050), cAMP (Sigma-Aldrich, D0627-1G) and doxycycline. Cells were differentiated in two steps: For pre-differentiation cycling (D00) cells were seeded at 2.5 x 10^6^ cells per T75 dish, grown for one day in proliferation medium. This was exchanged for differentiation medium for two days. On day 2 of differentiation cells were trypsinized, counted and seeded at 6 x 10^6^ per 10 cm dish, starting second step of differentiation. This is important because the differentiating neurons are very sensitive to cell density. During the subsequent differentiation process, half of the medium was changed every second day. Cells were taken for analysis, 6 (D06), 10 (D10) and 15 (D15) days after the initial exchange to differentiation medium.

### CRISPR

To delete *SNORD115* and *SNORD116* clusters, Alt-R CRISPR-Cas9 system from IDT was used: two crRNA:tracrRNA guides (for sequences see Supplementary Table 7) for upstream and downstream cleavage complexed with Alt-R Cas9 nuclease 3NLS (ThermoFisher Scientific, 1074182) were prepared using following protocol: 0.5 μl crRNA-U [200 μM], 0.5 μl crRNA-D [200 μM] and 1 μl tracrRNA-ATTO [200 μM] were annealed in the PCR machine: 5 min at 95°C, ramp -0.1°C/sec to 25°C. To deliver preassembled complexes and ssODN as a repair template, LUHMES cells were transfected by Nucleofection in a Nucleofector II device (Lonza) using a Basic Nucleofector Kit for primary neurons (Lonza, VAPI-1003) as described (Scholz *et al*., 2011). For each nucleofection reaction, the following proportions were used: 2 x 10^6^ cells, 100 μl nucleofector solution and 5μl of mix: 1.2 μl (120 pmol) crRNA:tracrRNA, 1.7 μl Cas9 (104 pmol), 0.5 μl ssODN [100 μM] and 1.6 μl PBS. 48 hours post nucleofection cells were FACS sorted into 96-well plate for the isolation of clones. Isolated clones were tested by PCR and by northern blot hybridization against SNORD115 and SNORD116 after neuronal differentiation. From CRISPR experiment we have obtained two heterozygous, paternal mutants for SNORD116 cluster (H116-1/1, H116-2/15) and only one mutant for SNORD115 cluster (H115-2/26). As the *SNHG14* gene is never expressed from maternal chromosome, we initially included one homozygous mutant (D115) in the analyses. However, this cell line subsequently gave quite different from all the heterozygotes at the level of transcriptome (in many cases appearing to be more similar to the wild-type). We therefore excluded it from further experiments and analyses used for final publication, other than the analyses of rRNA maturation.

### Northern blot

Depending on the size of the RNA of interest we used two different kinds of protocols. For snoRNAs: 10 μg total RNA was denatured in formamide loading dye and resolved on the 6% TBE-Urea gel (Novex, ThermoFisher Scientific) in 1 X TBE buffer, until the Bromophenol Blue dye left the gel. To verify even loading of the samples, the gel was stained with SYBRSafe (ThermoFisher Scientific) and scanned in FLA-5100 scanner (FujiFilm). RNA was transferred to the Nylon Hybond-N+ membrane (RPN303B; GE Healthcare) by wet electro-transfer using BioRad MiniProtean System, for 1 hour at 30 V. After the transfer, RNA was crosslinked to the membrane with UVC in Stratalinker. Prehybridization was done in UltraHyb-Oligo (ThermoFisher Scientific) for 2 hours at 42°C. Probes were hybridized overnight (5 pmol) in 15 ml UltraHyb-Oligo at 37°C. After washing, membrane was exposed on the storage phosphoscreen (BAS-MP2040, Fuji). After overnight exposure the screen was scanned in FLA-5100 scanner.

For rRNA: 2 µg total RNA was resolved on 1% agarose gel with TRI/TRI buffer and overnight capillary transfer onto BrightStar-Plus Positively Charged Nylon Membrane (ThermoFisher Scientific). The detailed method and hybridization conditions are published (Robertson *et al*, 2022). rRNA probe sequences were taken from: (Sloan *et al*, 2012; Tafforeau *et al*, 2013). A full list of probes can be found in Supplementary Table 7.

### RNA-seq libraries

For RNA-seq cells were grown on 10 cm dishes and lysed in 6 ml TRIZOL, frozen in two 3 ml aliquots. After phase separation, total RNA was collected in aqueous phase, mixed with 2 volumes of 100% ethanol, and further purified with Zymo Direct-Zol MiniPrep Plus kit (Zymo Research). 6 µg total RNA was treated with DNase RQ1 (Promega), purified with RNA Clean & Concentrator-5 kit (Zymo Research) and tested for integrity on Bioanalyzer using RNA 6000 Nano kit (Agilent). Ribosomal RNA was depleted from 0.8 μg total RNA using NEBNext rRNA Depletion Kit (New England Biolabs, E6350L) following the manufacturer’s protocol. RNA-seq libraries were prepared with NEBNext Ultra II Directional RNA Library Prep Kit for Illumina (New England Biolabs, E7765) and their quality was assessed on Bioanalyzer using High Sensitivity DNA assay (Agilent). RNA-seq libraries were sequenced by BGI Genomics. A list of samples and number of sequencings reads for each of them are provided in the Supplementary Table 1. This list includes also samples prepared from homozygous mutant D115. Although, surprisingly, qthis cell line is quite different from the remaining heterozygous cell lines, the observed differences may be meaningful and will potentially aid understanding to the role of the PWS cluster in the regulation of gene expression. We are therefore making the full datasets of samples publicly available.

### RNA-seq analysis

Sequencing reads preprocessed with **flexbar** (adapter trimming and quality filtering) (Dodt *et al*, 2012) were aligned to the genome (GRCh38 downloaded from Ensembl) with **STAR** (Dobin *et al*, 2013) (version=2.7.3a, `--outMultimapperOrder Random`) and aligned to the genomic features using **featureCounts** (version: 2.0.0, parameters: `-p -t exon -g gene_id -Q 10 - s 2`) (Dobin *et al*., 2013) and annotation from GENCODE (gencode.v34.annotation.gtf; evidence-based annotation of the human genome (GRCh38), version 34 (Ensembl 100) from 2020-03-24; (Harrow *et al*, 2012)). Differential expression analysis was performed using **EdgeR** package (Robinson *et al*, 2010) in RStudio (R Core Team; 2021). R: A language and environment for statistical computing. R Foundation for Statistical Computing, Vienna, Austria. https://www.R-project.org/). All the samples were combined in one DGEList, filtered by expression and normalized together, data dispersion was estimated with experimental design `(∼batch + group)` with group representing type of mutation at each timepoint e.g. WTD00, H115D15. Testing for differential expression was performed pairwise – mutation type vs WT strain for each differentiation time point with `glmTreat` function with recommended threshold of `lfc=log2(1.5)`. A list of differentially expressed genes is available in Supplementary Table 3. Functional enrichment analysis for differentially expressed genes (DEPs) between H116 and wild-type cells from all the differentiation stages combined was performed with g::Profiler (Raudvere *et al*, 2019) with the following parameters: data source: GO ontology: BP, MF CC; Statistical domain scope: all the genes included in EdgeR analysis, custom over all known genes; Significance threshold: g_SCS; user threshold: 0.05; electronic annotations [IEA] included. Most meaningful terms, selected manually with support from Revigo tool (Supek *et al*, 2011) are included in the plot created in RStudio with ggplot2 (cite: Wickham H (2016). ggplot2: Elegant Graphics for Data Analysis. Springer-Verlag New York. ISBN 978-3-319-24277-4, https://ggplot2.tidyverse.org).

PCA analysis was performed with prcomp function in RStudio using scaled log2(CPM) values and visualized with autoplot (Tang *et al*, 2016)

Analysis of differentiation was performed similarly to what is described above, with the following differences: only samples from wild-type cells were included in the analysis, data dispersion was estimated with experimental design `(∼ 0 + diffStage)`. Testing for differential expression was performed pairwise between the consecutive timepoints with `glmQLFTest` function with threshold `lfc=log2(2)`.

### Gene clustering

Clustering of genes based on the expression profile during differentiation was performed on the filtered by expression and normalized data from the EdgeR analysis of wild-type samples described above. Average expression for each gene (CPM) for given timepoint was calculated and normalized to the maximum expression for this gene, resulting in all the expression values falling in the range between 0 and 1. Those data were used as an input for k-means clustering with k=6 giving the best resolution without creating too much redundancy in the expression profiles. Functional enrichment analysis for genes belonging to each cluster was performed with g::Profiler tool (Raudvere *et al*, 2019), online version, with the following parameters: multiquery of 6 sets of genes CL0 to CL5; data source: GO ontology: Statistical domain scope: all the genes from CL0-CL5 clusters, custom over annotated genes; Significance threshold: g_SCS; user threshold: 0.05; electronic annotations [IEA] not included. List of the genes belonging in each cluster, excluding those filtered out by g::Profiler algorithm is provided in Supplementary Table 2, together with the detailed outcome of the analysis. Most meaningful terms, selected manually with some support from Revigo tool (Supek *et al*., 2011) are included in the plot created in RStudio with ggplot2 (Wickham, 2016) (https://ggplot2.tidyverse.org).

### FBL and FBLL1 alignment and structural analysis

Protein sequence alignment of FBL and FBLL1 was performed by Clustal Omega alignment tool (Larkin *et al*, 2007; Sievers *et al*, 2011) available within the SnapGene software (www.snapgene.com). Protein structure alignment was performed in PyMol (The PyMOL Molecular Graphics System, Version 2.5.4 Schrödinger, LLC) using AlphaFold structure predictions (Jumper *et al*, 2021; Varadi *et al*, 2022) of FBL and FBLL1 as input files.

### Alternative splicing analysis: DEXseq

STAR mapped RNA-seq data were aligned with **featureCounts** (Version 2.0.3) against flattened GTF file, i.e. genes with overlapping coordinates are combined into one composite gene (e.g. `gene_id` `"ENSG00000243485.5+ENSG00000284332.1").` Flattened file was produced by the `dexseq_prepare_annotation2.py` function downloaded from Github (Vivek Bharwaj, Subread_to_DEXSeq, Oct 27 2018 https://github.com/vivekbhr/Subread_to_DEXSeq) as recommended by DEXseq manual, with gencode.v34.annotation.gtf from GENCODE as an input file. As featureCounts doesn’t accept gene description longer than 256 bytes, 4 composite genes were removed from the analysis, among them *SNHG14* gene. featureCounts calculated reads mapping to exons and was run with the following parameters: `-p –countReadPairs -f -O -Q 10 -s 2`. Alternative splicing was analyzed with DEXSeq package (Anders *et al*., 2012) from Bioconductor project (Huber *et al*, 2015), independently for undifferentiated and d15 neurons, mutant vs WT cells, as well as for the differentiation of WT cells – WTD15_vs_WTD00, which was used as a positive control of the analysis.

### Calculating expression of SNHG14 exons

STAR mapped RNA-seq reads, were aligned with **featureCounts** against modified GTF file containing exclusively one type of features – exons, redundant exon annotations were removed (modified gencode.v34.annotation.gtf from GENCODE). Exon coverage data (CPM) were normalized to the size of the library and filtered using EdgeR package. Average values for an exon for each differentiation stage were calculated and normalized to the exon length. Distribution of normalized average expression values for exons within SNORD115 and SNORD116 clusters was visualized with ggplot2.

### GSEA/LEA analysis

Counts per million values (CPM) from EdgeR RNA-seq analysis (filtered and normalized to the size of the library) were used as an input for the Gene Set Enrichment Analysis (GSEA) (Subramanian *et al*., 2005). Experimental data were tested for enrichment in gene sets from Molecular Signature Database (MSigDB) (Subramanian *et al*., 2005), C8 collection: c8.all.v7.4.symbols.gmt using following parameters: permutation type: gene_set, number of permutations: 1000. For further analysis only gene sets originating from midbrain differentiation (MANNO_MIDBRAIN_PHENOTYPES) (La Manno *et al*., 2016) were followed. Genes that contribute to the distinct phenotypes were identified by Leading Edge Analysis (LEA, utility from GSEA). Functional enrichment analysis was performed on those genes using g::Profiler tool (Raudvere *et al*., 2019), online version, with the following parameters: multiquerry of 2 sets of genes WTd15-enriched and H116d15-enriched; data source: KEGG, TRANSFAC; Statistical domain scope: all the genes included in the GSEA analysis, custom over all known genes; Significance threshold: g_SCS; use threshold: 0.05; electronic annotations [IEA] not included. Detailed outcome of the analysis together with list of genes from LEA analysis is provided as Supplementary Table 6.

### MS samples

Samples were processed with modified FASP protocol (Wisniewski *et al*, 2009). Briefly, 0.5 x 10^6^ LUHMES cells (number of cells calculated at d2 of differentiation) were lysed in 100 μl lysis buffer (25 mM Tris-HCl pH 7.5, 50 mM DTT, 0.1% Rapigest), incubated in a thermoblock for 5 min at 95°C, mixing with 500 rpm, then allowed to cool to room temperature. Samples were sonicated at 4°C in Bioraptor Pico (Diagenode) for 10 cycles, 30 sec on, 30 sec off. 50 μl of the sample was mixed with 200 μl buffer B (8 M urea, 100 mM Tris-HCl pH 7.5), transferred onto the Vivacon 500 30k spin columns (Sartorius) and centrifuged at 14,000 x g for approximately 30 minutes until the buffer had all passed through. Proteins on the membrane were dissolved in 80 μl 100 mM iodoacetamide in 8 M urea and incubated in darkness for 20 min and centrifuged until the buffer had gone through. Samples were washed twice with 80 μl 50 mM ammonium bicarbonate (buffer ABC) and centrifuged until dry. 100 μl Trypsin solution (10 μg/ml in ABC) was added and samples are incubated overnight at 37°C. The next day peptide digest was collected by centrifugation into collection tube. Membrane is rinsed with 80 μl buffer ABC and both fractions were combined. Peptide concentration was measured on Qubit with Qubit Protein Assay (ThermoFisher Scientific) and samples were acidified by adding 10ul of 10% TFA. C18-stage tips were prepared as described (Rappsilber *et al*, 2003) and loaded with 10 μg tryptic peptides. StageTips, used to clean and concentrate the samples following digestion, were prepared as described (Rappsilber *et al*, 2007). Peptides were eluted in 40 μL of 80% acetonitrile in 0.1% TFA and concentrated down to 1 μL by vacuum centrifugation (Concentrator 5301, Eppendorf, UK). The peptide sample was then prepared for LC-MS/MS analysis by diluting it to 5 μL by 0.1% TFA.

LC-MS analyses were performed on an Orbitrap Fusion™ Lumos™480 Mass Spectrometer (Thermo Fisher Scientific, UK) coupled on-line, to an Ultimate 3000 HPLC (Dionex, Thermo Fisher Scientific, UK). Peptides were separated on a 50 cm (2 µm particle size) EASY-Spray column (Thermo Scientific, UK), which was assembled on an EASY-Spray source (Thermo Scientific, UK) and operated constantly at 50°C. Mobile phase A consisted of 0.1% formic acid in LC-MS grade water and mobile phase B consisted of 80% acetonitrile and 0.1% formic acid. Peptides were loaded onto the column at a flow rate of 0.3 μL min^-1^ and eluted at a flow rate of 0.25 μL min^-1^ according to the following gradient: 2 to 40% mobile phase B in 180 min and then to 95% in 11 min. Mobile phase B was retained at 95% for 5 min and returned to 2% a minute later, until the end of the run (220 min).

Survey scans were recorded at 120,000 resolution (scan range 350-1100 m/z) with an ion target of 8.0e5, and injection time of 50ms. MS2 Data Independent Acquisition (DIA) was performed in the orbitrap at 60,000 resolution, maximum injection time of 55ms and AGC target of 1.0E6 ions. We used HCD fragmentation (Olsen *et al*, 2007) with fixed collision energy of 30. From scan range 300-1000 m/z we used isolation windows of 17m/z and default charge state of 3. The desired minimum for points across the peak was set to 6.

The DIA-NN software platform (Demichev *et al*, 2020) version 1.8.1. was used to process the the DIA raw files and search was conducted against the Uniprot database (release of July 2017). Precursor ion generation was based on the chosen protein database (automatically generated spectral library from the protein database used) with deep-learning based spectra, retention time and IMs prediction. Digestion mode was set to specific with trypsin allowing maximum of one missed cleavage. Carbamidomethylation of cysteine was set as fixed modification. Oxidation of methionine, and acetylation of the N-terminus were set as variable modifications. The parameters for peptide length range, precursor charge range, precursor m/z range and fragment ion m/z range as well as other software parameters were used with their default values. The precursor FDR was set to 1%.

### MS data analysis

Differential expression of proteins was performed using DEP package (Zhang *et al*, 2018). Samples were filtered for proteins that are present in all replicates of at least one condition (`filter_missval(SE, thr = 0)`), VSN normalized and missing values were imputed with MinProb method (`q=0.01`) as most of the proteins are expected to be missing not at random (MNAR). Proteins are tested for differential expression using `test_diff` function and manually defined contrasts between conditions i.e. mutation type and stage of differentiation e.g. `H116d10_vs_WTd10`. Obtained p-values were corrected for multiple testing with Benjamini-Hochberg procedure using R stats package and all the proteins with p.adj ≤ 0.05 are treated as differentially expressed. A list of differentially expressed and all quantified proteins is available in Supplementary Table 5.

### Transcriptome-proteome analysis

Proteome (mean LFQ value for cell line at given differentiation stage e.g. PROT_Mean_WTD00) and transcriptome data (mean CPM value for cell line at given differentiation stage e.g. RNA_Mean_WTD00) after prior filtering and normalization steps described above, were combined in the same analysis.

Spearman correlations between steady state protein and RNA expression were calculated and ranged from 0.55 for undifferentiated cells to about 0.32 at D06 when the cells were still dynamically adjusting to changes connected with differentiation. Correlation values were dependent on the differentiation stage and very similar for all the cell lines (Fig. S6E). Despite moderate correlation of RNA-seq and MS data, clustering of combined datasets associated PROT and RNA data originating from the same cell type and differentiation stage (Fig. S6F). Because of the difference between LFQ and CPM values clustering was performed on the values scaled independently for PROT and RNA data (R, scale function). Clustering and plotting heatmap was performed with Pheatmap package (R, Version 1.0.12, Raivo Kolde, Pheatmap, 2019-01-04, https://CRAN.R-project.org/package=pheatmap). The ratio between protein and RNA expression (P/R) varies by the six orders of magnitude from ∼10^2^ up to ∼10^8^ (for tubulin TUBA4A). To test if this wide range of values reflects real conditions in the cells or just high noise in our sequencing data we compared P/R ratios for all genes, between cell lines at specific differentiation stages (P/R_MUT_)/(P/R_WT_) (Fig. S6). This value is quite stable as visible from a narrow distribution of values around 1 (full range of values: 0.003 to 81), suggesting that P/R ratio is a characteristic feature of a gene at a given differentiation stage and that can be utilized to test the hypothesis that ncRNAs from PWS locus influence post-transcriptional gene expression. We consider that direct influences of the ncRNAs on the stability of multiple proteins is unlikely.

We focused on genes that show changes in P/R ratio of at least two-fold upon deletion of the SNORD115 or SNORD116 cluster (Fig. S6I) for at least two differentiation stages and are identified as DEPs. This allowed us to limit the analysis to the most reliable subset of genes, with the protein expression stable enough to pass the statistical criteria of differential expression analysis. For all those genes we created a heatmap using ComplexHeatmap package (Gu *et al*, 2016).

### Author Contributions

AH and DT conceived the project and wrote the manuscript. AH and CS performed experiments. AH, TT, CS and DT analyzed data. All authors edited and reviewed the manuscript.

## Acknowledgements

We thank Shaun Webb for advice on bioinformatics, Tatsiana Auchynnikava for assistance with proteomics, Justyna Cholewa-Waclaw, Ruth Shah and Adrian Bird for the LUHMES cell line. We thank Dhanya Cheerambathur, Chris Sibley and colleagues for critical reading of the MS.

DT was supported by a Wellcome Principal Research Fellowship (077248). AH was supported by the Foundation for Prader-Willi Research (FPWR), TWT was supported by the Polish Ministry of Science and Higher Education Mobility Plus program (1069/MOB/2013/0). Work in the Wellcome Centre for Cell Biology is supported by a Centre Core grant (203149).

