## Supplementary material for "Roles of SNORD115 and SNORD116 ncRNA clusters in neuronal differentiation": Supp Figures 1 - 6

**Figure S1. Differentiation of LUHMES cells.**

A: Images of the differentiating wild-type LUHMES cells.

B: Heatmap of differentiating wild-type and mutant cells, including all RNA-seq samples with replicates.

**Figure S2. Construction of cell lines lacking SNORD115 of SNRORD116 expression.**

**A**: PCR confirmation of heterozygous deletion of SNORD115 (clone 2/12) and SNORD116 (clones 1/1 and 2/15) clusters in the cell lines involved in the current study. Homozygous SNORD115 deletion strain (D115) was used in the analysis of rRNA maturation only.

**B**: Time course of SNORD115 and SNORD116 expression in wild type and mutant cell lines.

**Figure S3. rRNA maturation is not affected by SNORD115 or DNORD116 deletion.**

**A**: Northern blots with probes against ITS1 (internal transcribed spacer 1), ITS2 and 5'ETS (external transcribed spacer) detecting rRNA maturation intermediates: 47/45S, 41S, 32S, 30S, 26S and 21S. Expression of mature 28S and 18S rRNA is visible on the scan of SybrSafe stained gel. Samples collected from wild-type and mutant cells, undifferentiated (D00) and at D10 of differentiation.

**Figure S4. Proteins forming intermediate filaments show strongly altered mRNA levels.**

A: Neurofilaments and associated protein mRNAs

Neurofilaments are generally composed of light, medium and heavy chains (NEFL, NEFM, and NEFH) and interact with other neuronal intermediate filament proteins Peripherin (PRPH) and Alpha-Internexin (INA). Nestin (Nes) and Vimentin (Vim) are unconventional neurofilament protein that promote neuronal morphogenesis [reviewed in (Bott & Winckler, 2020)].

B: Keratins and associated protein mRNAs.

**Figure S5. Deletion strains show minimal splicing defects, but altered transcription start and end sites.**

A: MPRIP gene changes splice variants upon differentiation.

B: Examples of genes identified by DEX-seq that display alternative start and termination sites, instead of expected alternative splice variants.

**Figure S6. Transcriptome-proteome analysis.**

A: Pearson correlation plot of mass spectrometry samples.

B: Expression of all quantified proteins across differentiation process in wild-type cells.

C: Pattern of expression of all identified differentially expressed proteins (DEPs) in wild-type and mutant cells.

D: Number of differentially expressed proteins in each mutant cell line increases with the course of differentiation.

E: Proteome and transcriptome samples cluster together. Clustering performed on scaled data.

F: Relation between transcript and protein levels in differentiating LUHMES cells. Spearman correlation ($\rho$) is higher in cycling than differentiating cells.

G: The steady-state amount of protein per steady-state amount of transcript (P/R ratio) is highly variable.

H: P/R ratio is a characteristic feature of a gene at a given differentiation stage and is generally maintained between wild-type and mutant cells. This plot does not contain all values; outliers were removed to Altered the narrow range of values around 1.

I: Correlation of P/R ratios between wt and H116 during differentiation. Higher and lower values were observed in the mutant. $\rho$, Spearman correlation.

**Figure S7**. Representative enrichment plots from analyses comparing LUHMES cells and cells from differentiating human midbrain

A: Enrichment plots for cell types with similar transcriptional profile to H116 deletion mutant; HDA2 – dopaminergic neurons, subtype 2; HSERT – serotonergic neurons; HNBML5 - mediolateral neuroblasts, subtype 5.

B: Enrichment plots for cell types with similar transcriptional profile to wild-type LUHMES cells at day 15 of differentiation; HPROGBP – progenitor basal plate; HRGL2B – radial glia-like cells, subtype 2B. Midbrain data from (La Manno *et al.*, 2016) deposited in the Molecular Signatures Database (MSigDB) (Subramanian *et al.*, 2005).

**SUPPLEMENTARY TABLES: 1-7**

**Supplementary Table 1**

List of the RNAseq and Mass Spectrometry samples

**Supplementary Table 2**

Detailed outcome of the GO term analysis of clusters.

**Supplementary Table 3**

Differentially expressed genes from RNA-seq data analysis.

**Supplementary Table 4**

Detailed outcome of the GO term analysis of DEGs between H116 and wild type cells.

**Supplementary Table 5**

Differentially expressed proteins from Mass Spectrometry data analysis.

**Supplementary Table 6**

Transcription factors and KEGG pathway analysis of genes associated with “mature” phenotype of H116 cells.

**Supplementary Table 7**

List of oligonucleotides for CRISPR, PCR and Northern blot.

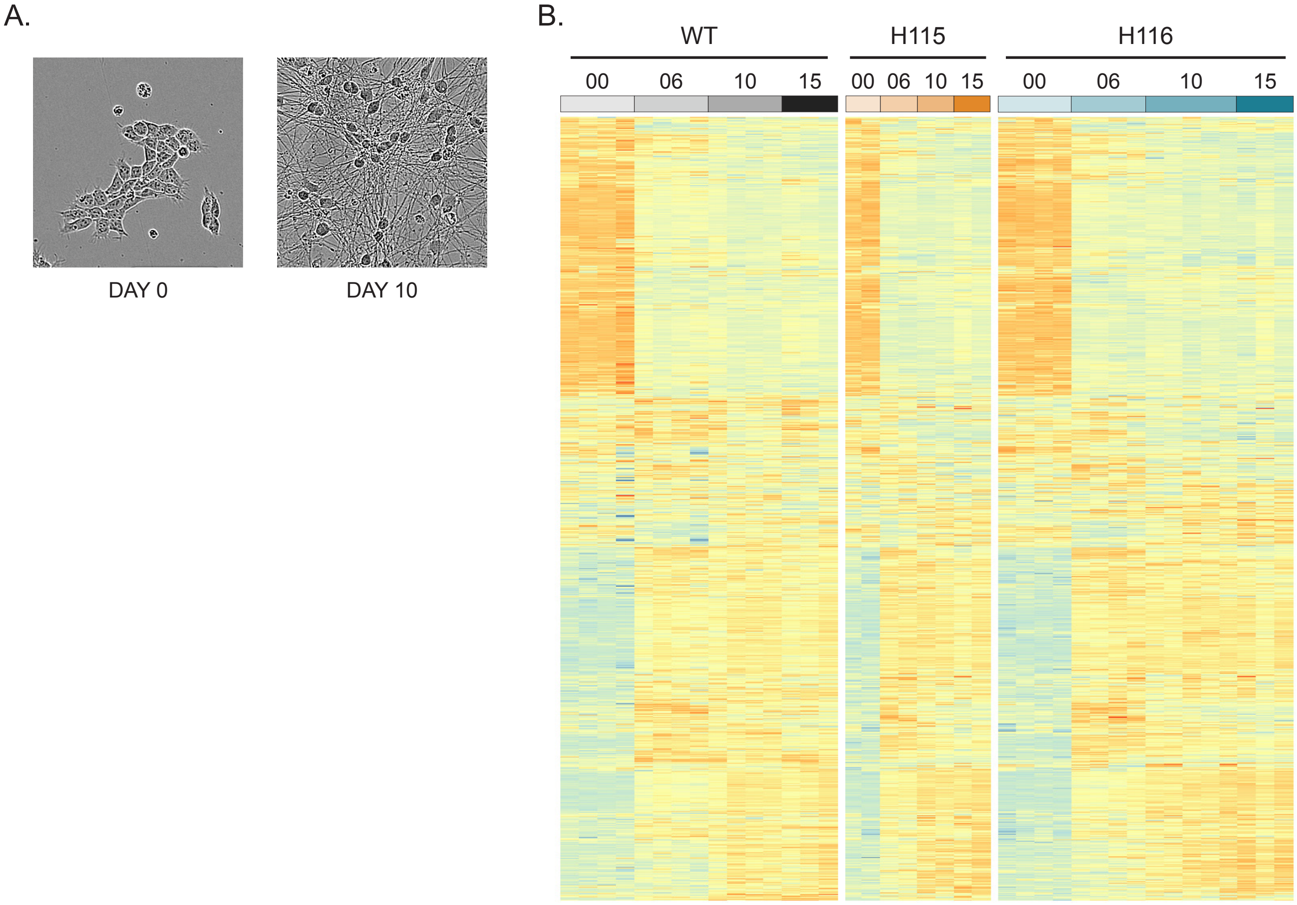


**Fig. S1**

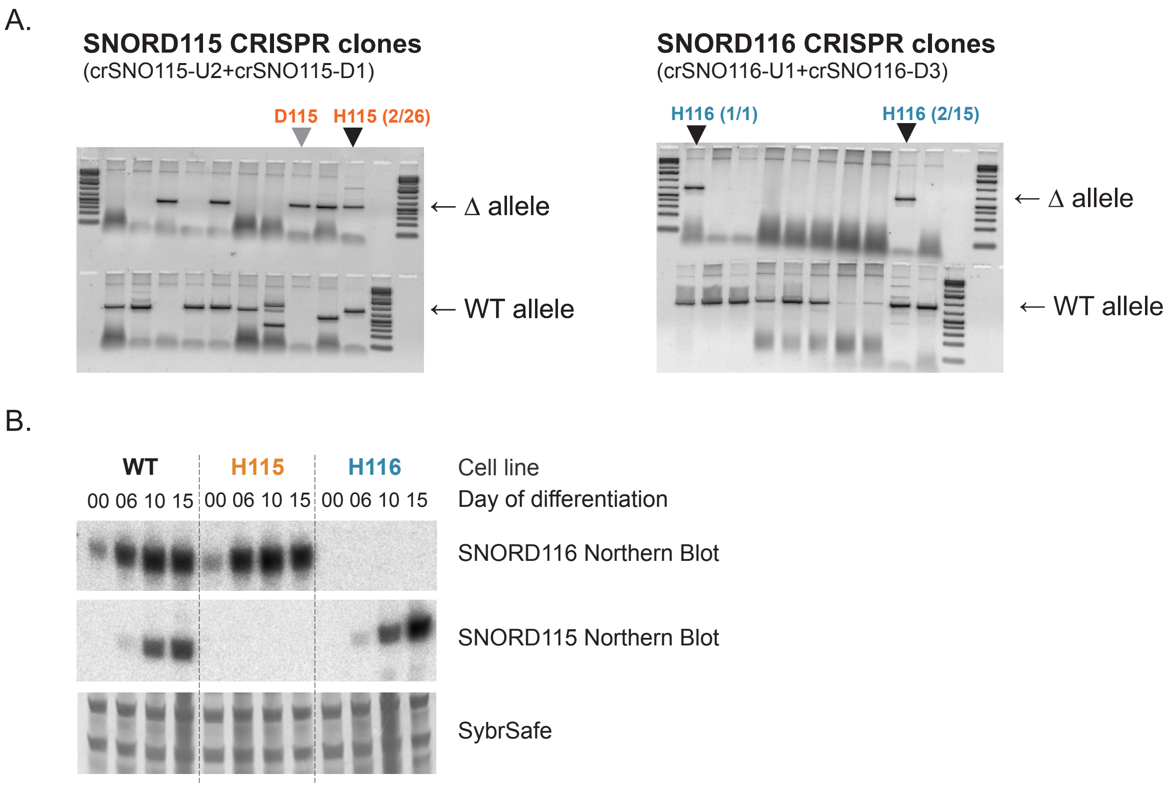
A.
SNORD115 CRISPR clones
(crSNO115-U2+crSNO115-D1)
D115
H115 (2/26)
← Δ allele
← WT allele
SNORD116 CRISPR clones
(crSNO116-U1+crSNO116-D3)
H116 (1/1)
H116 (2/15)
← Δ allele
← WT allele
B.
WT
H115
H116
Cell line
00 06 10 15 00 06 10 15 00 06 10 15
Day of differentiation
SNORD116 Northern Blot
SNORD115 Northern Blot
SybrSafe

### Fig. S2

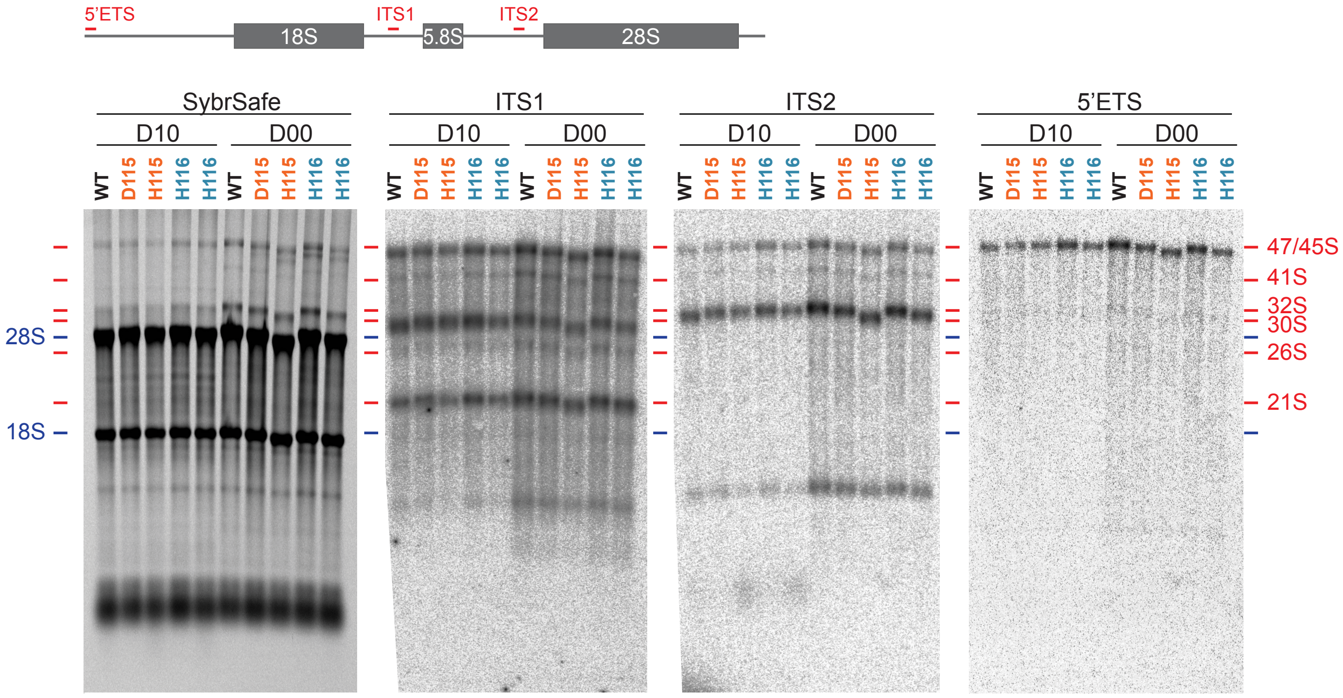
5'ETS
18S
ITS1
5.8S
ITS2
28S
SybrSafe
ITS1
ITS2
5'ETS
D10
D00
WT
D115
H115
H116
28S
18S
47/45S
41S
32S
30S
26S
21S

**Fig S3.**

A.

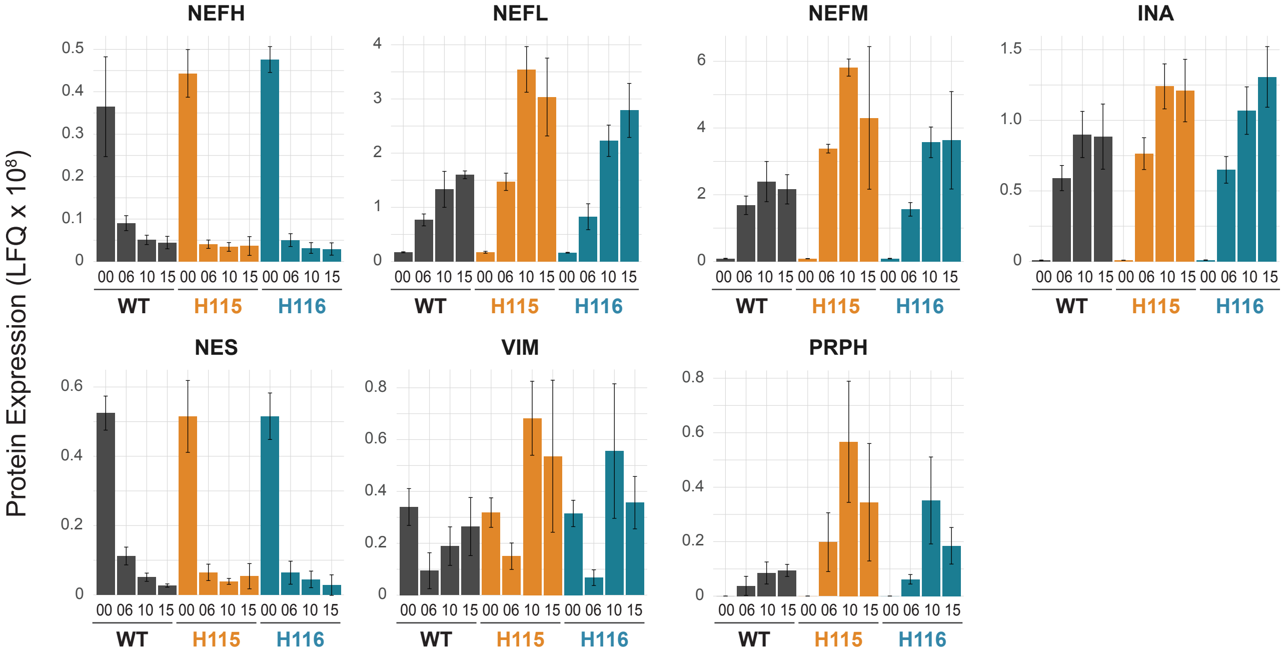

NEFH
NEFL
NEFM
INA
NES
VIM
PRPH
Protein Expression (LFQ x 10^8)
00 06 10 15 00 06 10 15 00 06 10 15
WT
H115
H116


B.

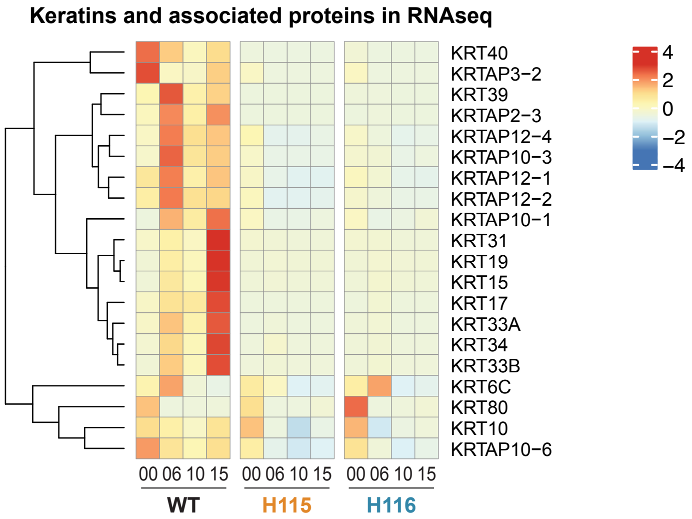

Keratins and associated proteins in RNAseq
KRT40
KRTAP3−2
KRT39
KRTAP2−3
KRTAP12−4
KRTAP10−3
KRTAP12−1
KRTAP12−2
KRTAP10−1
KRT31
KRT19
KRT15
KRT17
KRT33A
KRT34
KRT33B
KRT6C
KRT80
KRT10
KRTAP10−6
4
2
0
−2
−4
00 06 10 15 00 06 10 15 00 06 10 15
WT
H115
H116


### Fig. S4

A.

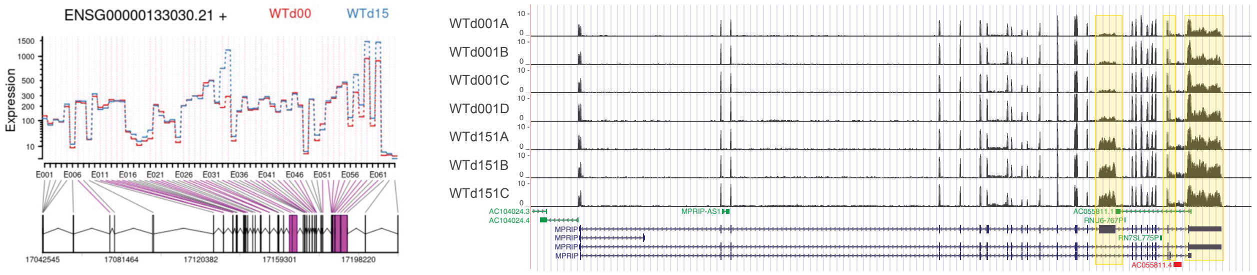
ENSG00000133030.21 +
WTd00
WTd15
Expression
WTd001A
WTd001B
WTd001C
WTd001D
WTd151A
WTd151B
WTd151C
MPRIP

B.

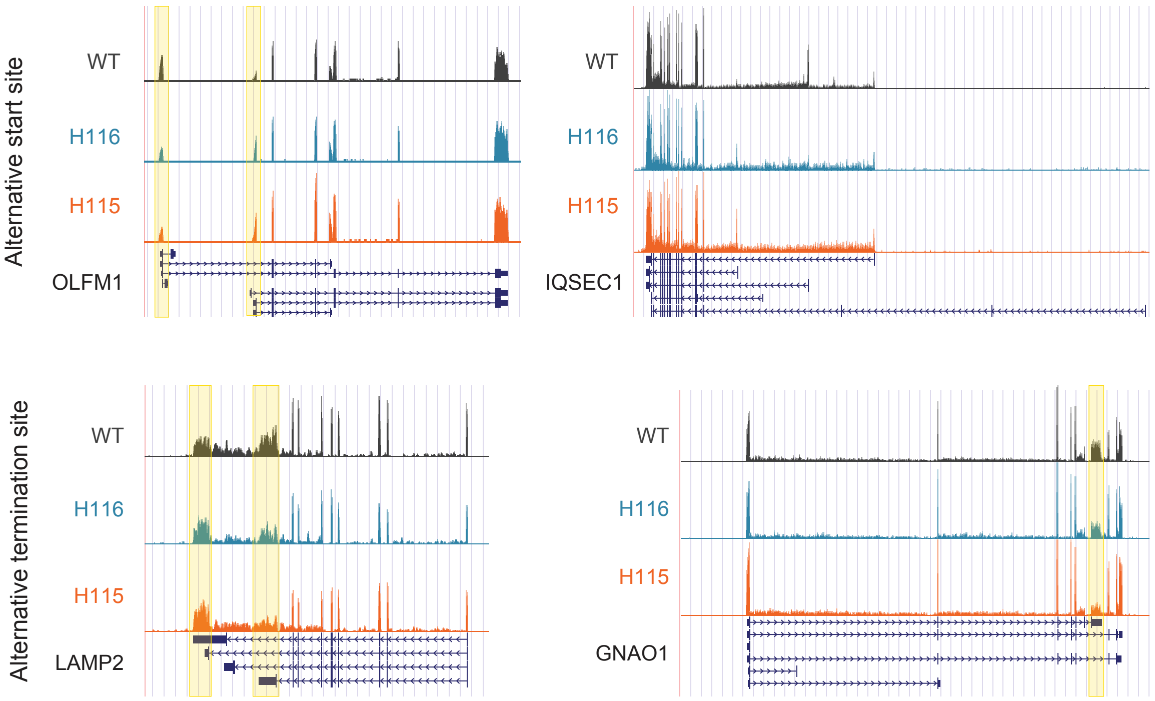
Alternative start site
WT
H116
H115
OLFM1
WT
H116
H115
IQSEC1
Alternative termination site
WT
H116
H115
LAMP2
WT
H116
H115
GNAO1

Fig. S5

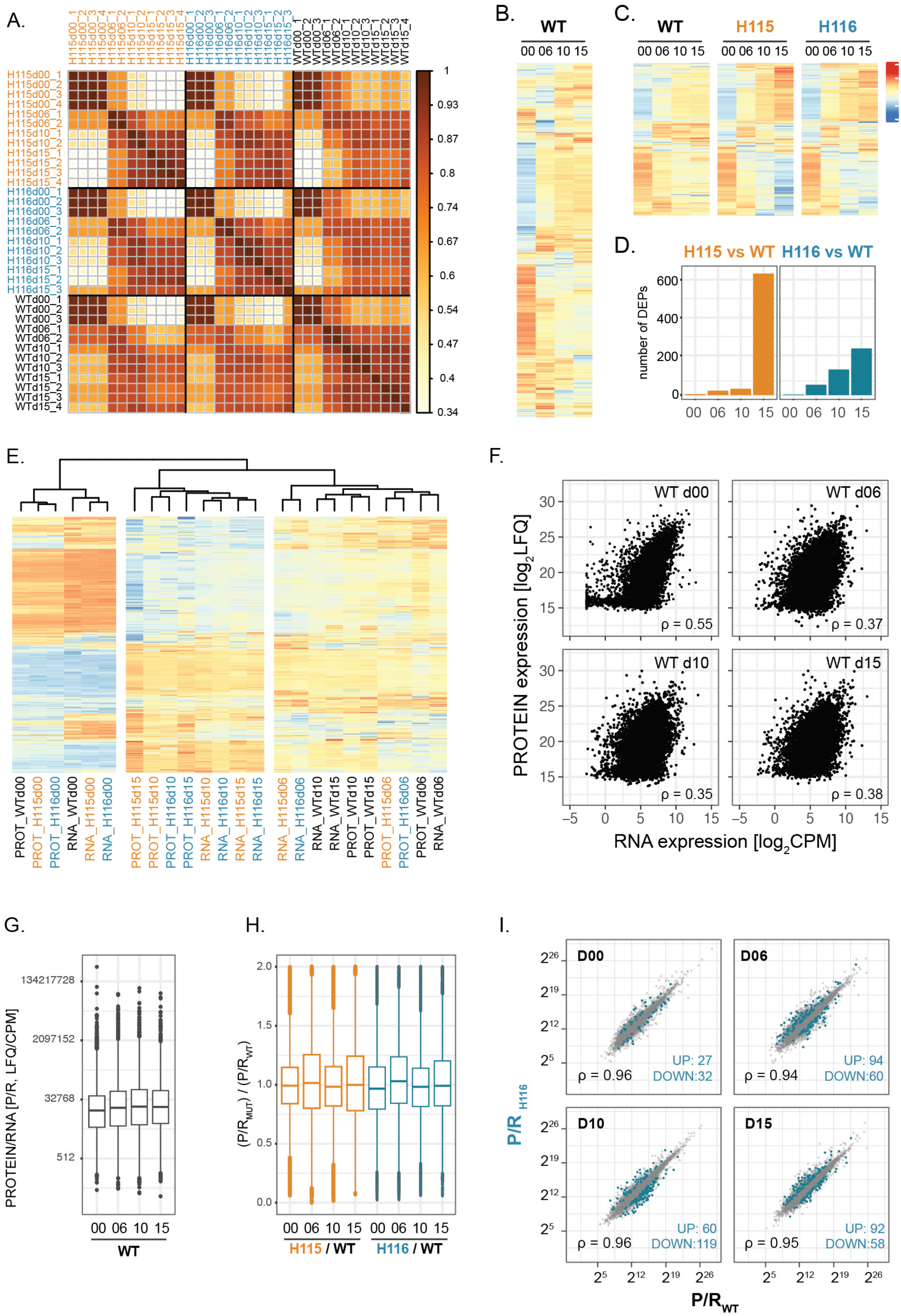
A.
B.
WT
00 06 10 15
C.
WT
H115
H116
D.
H115 vs WT
H116 vs WT
number of DEPs
E.
F.
WT d00
WT d06
WT d10
WT d15
ρ = 0.55
ρ = 0.37
ρ = 0.35
ρ = 0.38
PROTEIN expression [log2LFQ]
RNA expression [log2CPM]
G.
PROTEIN/RNA [P/R, LFQ/CPM]
H.
H115 / WT
H116 / WT
I.
D00
D06
D10
D15
ρ = 0.96
UP: 27
DOWN:32
ρ = 0.94
UP: 94
DOWN:60
ρ = 0.96
UP: 60
DOWN:119
ρ = 0.95
UP: 92
DOWN:58
P/R H116
P/R WT

**Fig S6.**

A.

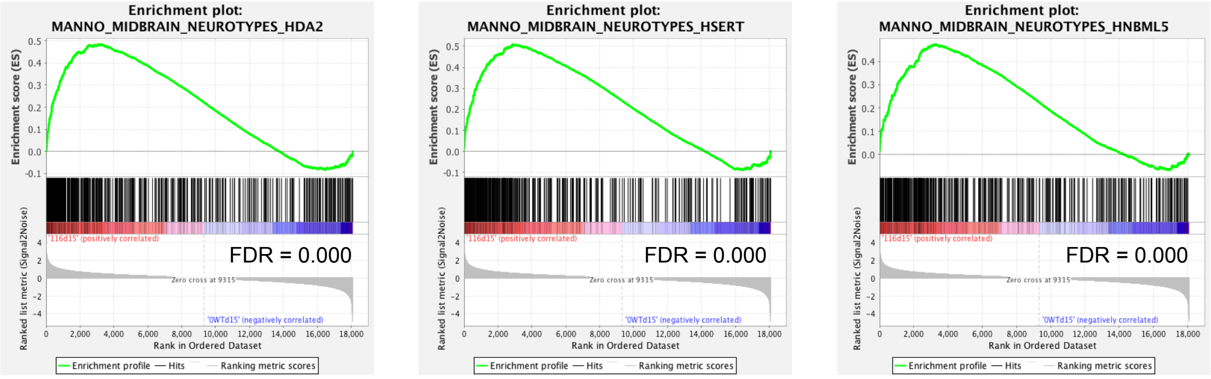
Enrichment plot:
MANNO_MIDBRAIN_NEUROTYPES_HDA2
FDR = 0.000
Enrichment plot:
MANNO_MIDBRAIN_NEUROTYPES_HSERT
FDR = 0.000
Enrichment plot:
MANNO_MIDBRAIN_NEUROTYPES_HNBML5
FDR = 0.000
Enrichment score (ES)
Ranked list metric (Signal2Noise)
'116d15' (positively correlated)
Zero cross at 9315
'0WTd15' (negatively correlated)
Rank in Ordered Dataset
Enrichment profile
Hits
Ranking metric scores

B.

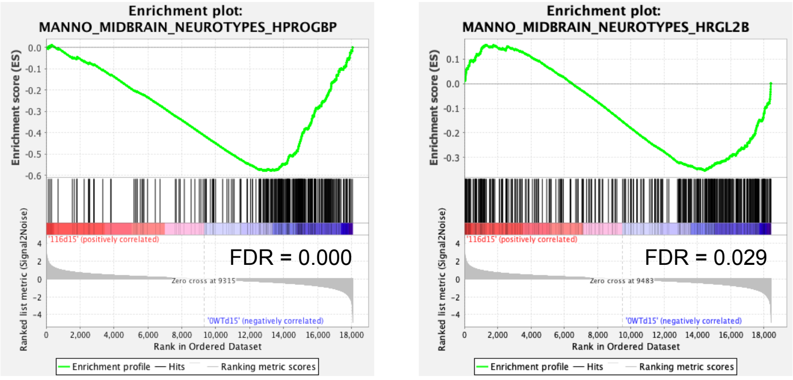
Enrichment plot:
MANNO_MIDBRAIN_NEUROTYPES_HPROGBP
FDR = 0.000
Zero cross at 9315
Enrichment plot:
MANNO_MIDBRAIN_NEUROTYPES_HRGL2B
FDR = 0.029
Zero cross at 9483
Enrichment score (ES)
Ranked list metric (Signal2Noise)
'116d15' (positively correlated)
'0WTd15' (negatively correlated)
Rank in Ordered Dataset
Enrichment profile
Hits
Ranking metric scores

**Fig. S7**
